# High piRNA abundance is a hallmark of evolutionarily persistent piRNA-producing loci

**DOI:** 10.64898/2026.11.04.764267

**Authors:** Adrià Mitjavila-Ventura, Samuele M. Amante, Cristina Moreta-Moraleda, Sonia V. Forcales, Miguel R. Branco, Tanya Vavouri

## Abstract

PIWI-interacting RNAs (piRNAs) silence transposons in the germline, and the loci that produce them diverge rapidly between species. Yet, some loci retain both their genomic location and their piRNA production across large evolutionary distances, and the basis for this conservation is unclear. To identify the features that distinguish these persistent loci, we compared piRNA production at orthologous loci in three closely related mouse species, contrasting loci that produce piRNAs in all three species with those that do so in only one. Transposon content and UTR features, often linked to piRNA production, did not distinguish the two groups. Transcription extending beyond gene ends (readthrough) was more frequent at conserved loci, but did not explain conservation once the amount of piRNAs produced was taken into account. Instead, we found that the conserved loci are those that produce the most piRNAs. The same association holds true in human and two other primate species. We speculate that most piRNAs expressed in testis are tolerated molecular noise, and that only the highly expressed minority are likely to be functional.

## BACKGROUND

PIWI-interacting RNAs (piRNAs) are small non-coding RNAs that guide PIWI proteins to silence transposable elements (TEs) and regulate the expression of certain genes in the animal germline. In doing so, piRNAs safeguard genome integrity and contribute to proper gene regulation during gametogenesis [1–4]. In most animals, piRNAs are processed from long single-stranded precursors transcribed from discrete coding and non-coding loci, termed piRNA-producing loci, or piRNA clusters. Although the piRNA pathway is conserved in nearly all animals [5,6], the loci producing piRNAs and especially piRNAs themselves are remarkably different. The sequence of piRNA precursor transcripts evolves under weak selective constraint [7–11], and even conserved piRNA-producing loci may produce different piRNAs and target distinct transcripts in different species. With few exceptions, piRNAs and piRNA clusters vary greatly between species [10,12]. Indeed, many piRNA clusters are species-specific, even when overlapping conserved protein-coding genes [7,10].

Despite the widespread conservation of the piRNA pathway, few piRNA-producing loci have been proven to have a function [13,14]. This raises the possibility that many of these loci may be producing piRNAs with little or no benefit to the host and evolving neutrally [15]. Comparative studies of piRNAs from different individuals from the same species of fly [12,16], zebrafish [17], and mouse [18] have shown that within a species there are loci with polymorphic piRNA expression explained by genetic variation. Extending this comparative analysis across species can reveal properties associated with evolutionary persistence and thus offer insights into the properties conferring fitness benefit to the host.

To address this, we compared piRNA clusters in three closely related mouse species, the house mouse *Mus musculus*, the Ryukyu mouse *Mus caroli*, and the Gairdner’s shrew mouse *Mus pahari* (**fig. 1A**). We chose these species because they are closely related enough that many clusters remain conserved, yet sufficiently diverged to reveal differences in piRNA production, while having genomes that are highly alignable [19]. By comparing loci that produce piRNAs in all three species with those that are species-specific, we tested how different sequence and transcript properties, including transcript expression, piRNA abundance, transposon content, UTR length and transcriptional readthrough, relate to the evolutionary persistence of piRNA clusters.

**Figure 1.**
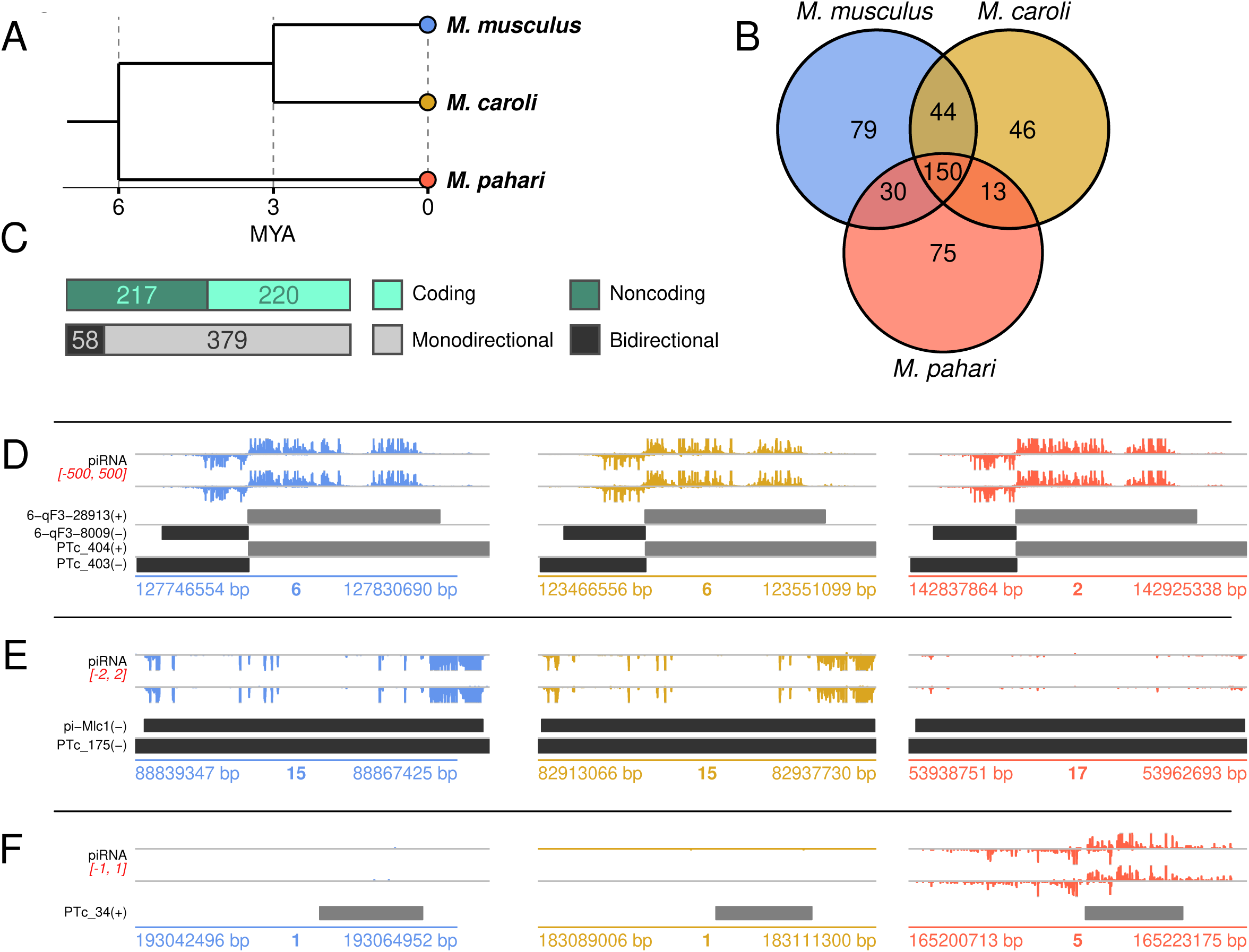
Divergence of piRNA production from orthologous loci in three closely-related mouse species. (A) Phylogenetic tree of Mus musculus, Mus caroli and Mus pahari, with evolutionary distances retrieved from Thybert et al. [19]. (B) Species in which the 437 piRNA-producing loci are expressed. (C) Proportion of piRNA clusters overlapping protein-coding genes (upper barplot) and proportion of piRNA clusters annotated as monodirectional or bidirectional by a custom function (lower barplot). (D-F) Genomic tracks of three predicted piRNA clusters in each of the species: (D) PTc_403 and PTc_404 are two conserved piRNA clusters producing piRNAs in all three species; (E) PTc_175 is a locus producing piRNAs only in *M. musculus* and *M. caroli*; and (F) PTc_34 is a locus producing piRNAs only in *M. pahari*.

If piRNA production from a locus were a consequence of a property of the locus itself, such as the length of its 3’ UTR, then that property, rather than the amount of piRNAs produced, should predict conservation. If instead the piRNAs produced from a locus are themselves selected, for example because they regulate other transcripts *in trans*, then conservation should correlate with the number and abundance of piRNAs the locus produces. Here we test these predictions in three mouse species and then in primates. We find that piRNA output is the strongest predictor of conservation. This points to high piRNA output as a way to identify, among the many loci that produce piRNAs, the few whose piRNAs are likely to benefit the host.

## RESULTS

### 1. Nearly half of mouse piRNA clusters are less than six million years old

We started by defining the loci that produce piRNAs in each mouse species and quantifying their piRNA expression. For each species, including *M. musculus*, we sequenced small RNAs from testes of two young adults (**table S1**). All samples showed typical characteristics of piRNAs [4]; small RNA length between 24 and 32 nucleotides with a mode length of 30nt (**fig. S1A**) [1], strong bias for uridine in the first nucleotide (**fig. S1B**) [2,3], enrichment of antisense pairs overlapping by 10nt at the 5’ end (**fig. S1C**) [20], and enrichment of stop codons in the first three nucleotides (**fig. S1D**) [21]. We then aligned all small RNAs to the genome of the corresponding species and predicted piRNA clusters, retaining only those predicted in both samples from each species (**fig. S1E**, **table S2**).

To assess the quality of the predicted piRNA clusters, we compared the 311 *M. musculus* piRNA clusters predicted from this dataset (**table S2A**) with the high-confidence mouse piRNA clusters (also from *M. musculus*) previously defined using multiple sources of evidence [22,23] (referred to as Yu2021 piRNA clusters, **table S3A**). The predicted piRNA clusters from this study show a similar ratio of divergently transcribed (i.e. bidirectional) versus monodirectionally transcribed piRNA clusters, and a similar ratio of piRNA clusters overlapping protein-coding genes with the Yu2021 piRNA clusters (**fig. S1F**). Of the 311 predicted piRNA clusters in *M. musculus*, 121 overlapped Yu2021 piRNA clusters (**fig. S1G**), and among the remaining 190 loci, 162 (∼85.3%) intersected with piRNA clusters predicted from an independent set of five inbred mouse strains [18] (**fig. S1H**). Of the 121 predicted loci overlapping Yu2021 piRNA clusters, 71 are piRNA clusters defined as pachytene according to their expression pattern during spermatogenesis (**fig. S1I**), reflecting the overrepresentation of pachytene and post-meiotic germ cells in whole testes of adult mice [22,24]. Conversely, of the 94 Yu2021 piRNA clusters not overlapping piRNA clusters predicted here, 53 piRNA clusters (56%) are expressed selectively at pre-pachytene stages of sperm development (**fig. S1I**) and therefore their underrepresentation in this dataset is expected. For simplicity, we refer to the predicted piRNA clusters as piRNA clusters and the small RNAs that map to them as piRNAs throughout the rest of the manuscript. For interspecies comparisons, we converted the coordinates of all piRNA clusters from all three species to the same coordinate system, merged overlapping loci and removed those in unalignable or duplicated regions. Finally, using an expression threshold we defined 437 orthologous loci that produce piRNAs in one, two, or three species (see ‘**Methods**’). Hereon, we refer to a locus as a piRNA cluster in the species in which piRNA abundance is above the predefined threshold; each of the 437 loci is defined as a piRNA cluster in one, two, or all three species (**fig. 1B**, **fig. S1E**, **table S3B**). From this final set of loci, in *M. musculus*, half overlap protein-coding genes and most produce piRNAs from only one strand (**fig. 1C**). As expected, piRNA expression is higher in the species in which the locus is defined as a piRNA cluster (**fig. S1J**). Of the 437 loci, 200 (∼46%) are species-specific piRNA clusters, therefore likely gained within the last six million years, 44 (∼10%) are shared only between the two closely related mouse species (*M*. *musculus* and *M. caroli*), 43 (∼10%) clusters are either ancestral and lost in one species, or independently gained in two of the three species *(M. musculus* and *M. pahari* or *M. caroli* and *M. pahari*), and 150 (∼34%) are conserved in all three species (**fig. 1B,D-F**). The comparison of three closely related species highlights the short-lived nature of most mouse piRNA clusters. We used this set to ask which genomic features contribute to the persistence of piRNA-producing loci.

### 2. Transposon content does not relate to conservation of piRNA production

We and others previously reported that polymorphic piRNA-producing loci are enriched in polymorphic TE insertions, particularly endogenous retroviruses (ERVs) [12,16,18]. In this dataset we also detect species-specific piRNA clusters that co-occur with TEs annotated in the same species (**fig. S2**), such as the *A1cf* locus (cluster PTc_244), which contains a LINE1 element flanked by two IAP solo LTRs and shows increased mRNA and piRNA expression in *M. musculus* (**fig. S2A**); or the locus antisense to *Ptprm* (cluster PTc_225) and a locus on chromosome 3qA3 between *Kcnmb3* and *Zfp639* (cluster PTc_294), both of which harbour a RLTR10B insertion near their promoter and produce piRNAs only in *M. musculus* (**fig. S2B,C**).

To test for the enrichment of individual TE families in piRNA clusters in a systematic way, we compared transposon insertion frequencies in piRNA clusters against 1000 permutations in which piRNA cluster positions were randomised (**fig. 2A**, **table S4)**. We found that most TE families are depleted in piRNA clusters compared to random genomic regions, as previously reported [25] (**fig. 2A)**. The exceptions are mostly LTRs, which are observed more than expected by chance (**fig. 2A**). Compared to expression-matched genes, general TE content for the three major retrotransposon classes is significantly higher in piRNA clusters (**fig. 2B**, **table S5**). Therefore, piRNA clusters have higher tolerance for TE insertions than genes but lower tolerance compared to the rest of the genome. Finally, we asked whether species-specific and conserved piRNA clusters differ in their general TE content, but found no significant differences (**fig. 2C**, **table S5C**). Thus, general TE content does not predict the conservation of piRNA clusters.

**Figure 2.**
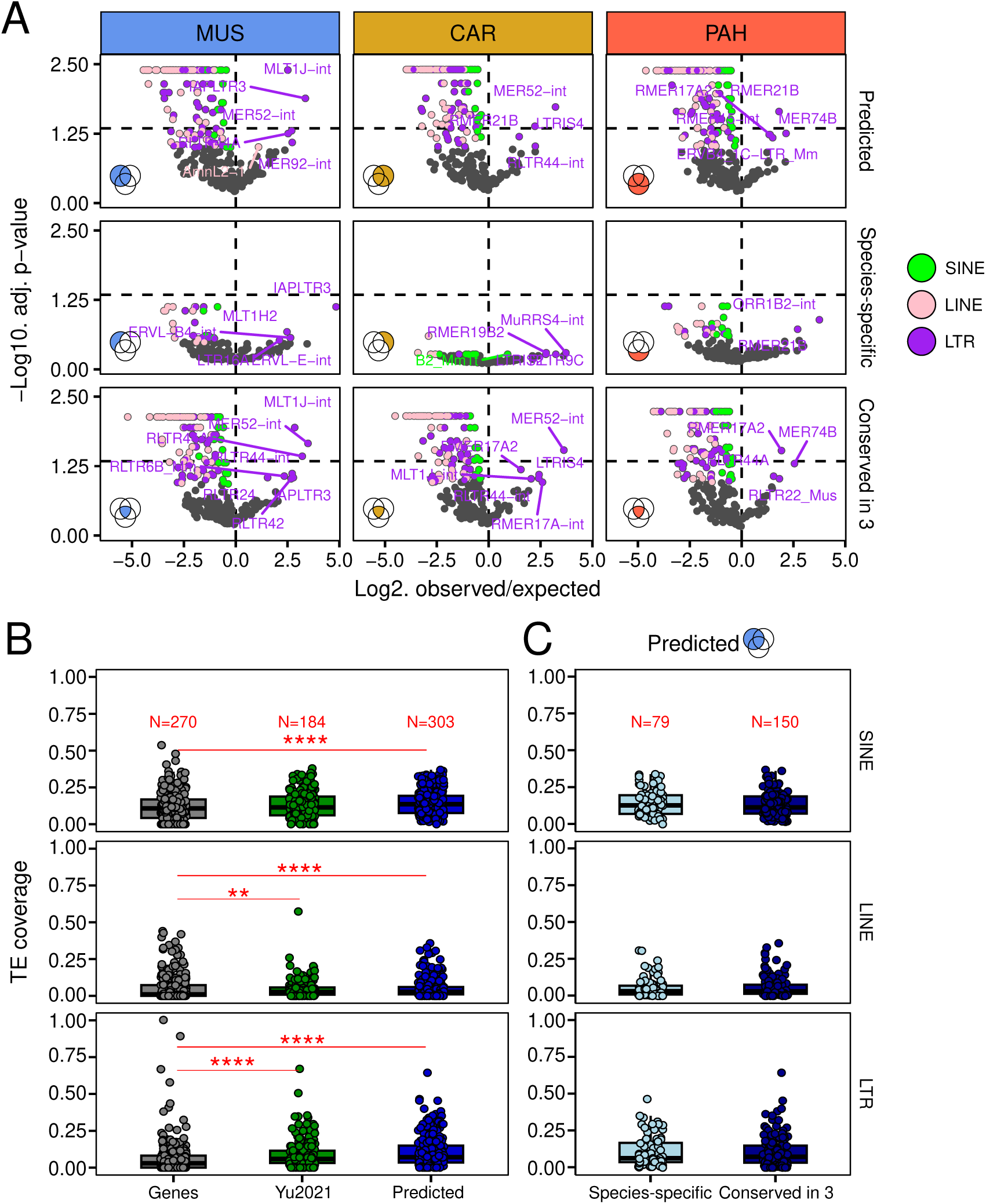
Transposon content in predicted piRNA clusters does not relate to piRNA cluster conservation. (A) TE families enriched and depleted in predicted piRNA clusters, according to number of overlaps in 1000 permutations performed with *regioneReloaded* R package. Dots are coloured by TE class when nominally significant (p<0.05) and grey otherwise; the dashed line marks the stricter Benjamini-Hochberg adjusted p-value threshold of 0.05. Rows show, from top to bottom, all predicted clusters in a species, species-specific clusters, and clusters conserved in all three species. (B) TE coverage in Yu2021 (N=184) and predicted (N=303) piRNA clusters compared to expression-matched control genes (N=270). (C) TE coverage in species-specific (N=79) and conserved (i.e., Conserved in all three species, N=150) piRNA clusters. Significance values in panels B and C are from Wilcoxon rank sum (non paired) test. Significance codes: *: p<0.05, **: p<0.01, ***: p<0.001, ****: p<0.0001.

### 3. piRNA-producing genes have longer 3’ UTRs but UTR features do not predict cluster **conservation**

In many animals, including flies, frogs, and mice, a subset of germline-expressed protein-coding genes produce piRNAs from their 3′ untranslated regions (UTRs) [26,27]. As with other piRNA-producing loci, a fraction of these genic piRNA clusters is conserved across species **(e.g. fig. 3A),** while many others are species-specific (e.g. **fig. 3B**), raising the question of whether properties of UTRs influence their evolutionary persistence.

**Figure 3.**
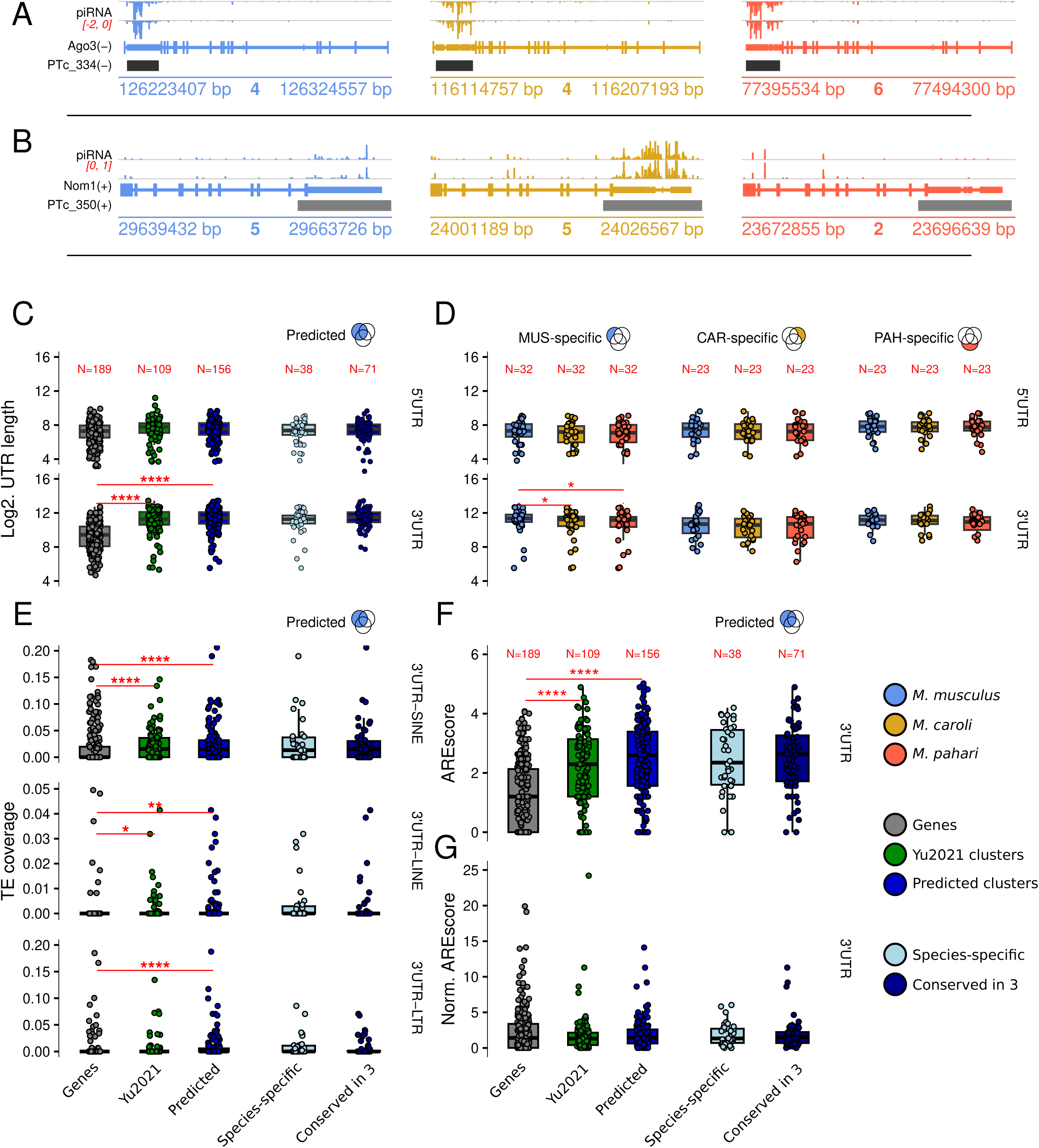
piRNA-producing genes have longer 3’ UTRs, but UTR features do not predict piRNA cluster conservation. (A,B) Genomic tracks of conserved (A) and species-specific (B) piRNA clusters overlapping 3’UTRs. (C) 5’ and 3’ UTR length (top and bottom rows) of (left) genic Yu2021 (N=109) and predicted (N=156) piRNA clusters compared to expression-matched control protein-coding genes (N=189) and (right) species-specific (N=38) versus conserved (N=71) piRNA clusters. (D) 3’UTR length of protein-coding species-specific piRNA clusters and their non piRNA-producing orthologs in the other species (N(MUS)=32, N(CAR)=23 and N(PAH)=23). (E) TE coverage in 3’UTRs of (left) coding piRNA clusters (N(Yu2021)=109 and N(predicted)=157) compared to control genes (N=189) and (right) species-specific vs conserved piRNA clusters. Rows are SINE, LINE and LTR. (F) AU-rich element (ARE) enrichment in 3’UTRs of (left) piRNA clusters compared to control genes (N=189) and (right) species-specific versus conserved piRNA clusters. ARE enrichment is measured using a custom implementation of AREscore. (G) AREscore normalized by length in 3’UTRs of (left) piRNA clusters compared to control genes (N=189) and (right) species-specific versus conserved piRNA clusters. Significance values in panels C, E, F and G are from Wilcoxon rank sum (non paired) test. Significance values in panel D are from Wilcoxon signed rank (paired) test. Significance codes: *: p<0.05, **: p<0.01, ***: p<0.001, ****: p<0.0001.

We first asked whether UTR length correlates with piRNA production. Annotated 3’ UTRs of piRNA-producing genes from *Yu et al. (2021)* [23] and the ones defined in this study are significantly longer than those of expression-matched control genes, while 5’ UTRs are not (**fig. 3C, table S6**). However, conserved and species-specific genic piRNA clusters do not significantly differ in UTR length (**fig. 3C**). Furthermore, species-specific genic piRNA clusters and their non piRNA-producing orthologous loci do not differ in UTR length (**fig. 3D, table S6D**), except for species-specific clusters from *M. musculus.* The difference in this comparison most likely reflects the more complete 3′ UTR annotation available for *M. musculus,* whose reference-quality genome is supported by far more transcriptomic evidence than those of *M. caroli* and *M. pahari,* rather than a genuine biological difference (**fig. S3**). Overall, 3’ UTR length is a feature of genes that produce piRNAs but does not distinguish orthologs that produce piRNAs from those that do not, nor does it predict whether that production is conserved.

We then tested whether TEs in 3’ UTRs are linked to piRNA production. We found higher SINE (and other TE) coverage in 3′ UTRs of piRNA-producing genes relative to non-piRNA-producing genes, but no significant difference between conserved and species-specific piRNA clusters (**fig. 3E, table S7**). Protein-coding genes that produce piRNAs may have a higher tolerance for TEs in their 3’ UTRs but the presence of TEs has no detectable effect on the conservation of piRNA production from these genes.

Long UTRs could also influence piRNA production by harboring additional regulatory elements. For instance, conserved 3′ UTR piRNA clusters in mouse have been reported to harbour structured RNA elements and poly-U-rich motifs [10]. Therefore, we tested whether AU-rich elements (AREs) are associated with persistence of piRNA production. The 3’ UTRs of piRNA-producing genes are significantly enriched in AREs relative to control genes (**fig. 3F, table S8**), but this enrichment is lost after controlling for UTR length (**fig. 3G, table S8**). Moreover, conserved and species-specific piRNA clusters do not differ in ARE content (**fig. 3F-G**). Taken together, these results indicate that while piRNA-producing genes have longer UTRs, neither UTR length, TE content, nor ARE content predicts the conservation of piRNA production.

### 4. Transcriptional readthrough is more common among conserved piRNA clusters

piRNAs are not only produced from annotated 3’ UTRs of genes but often also extend downstream (examples shown in **fig. 4A,B**). This phenomenon, known as transcriptional readthrough or transcription downstream-of-gene (DoG), occurs when transcription continues beyond the canonical transcription termination site [28]. Recent studies have reported transcriptional readthrough as a mechanism for piRNA production in several species [29,30]. We therefore investigated the relationship between transcriptional readthrough and piRNA cluster conservation.

**Figure 4.**
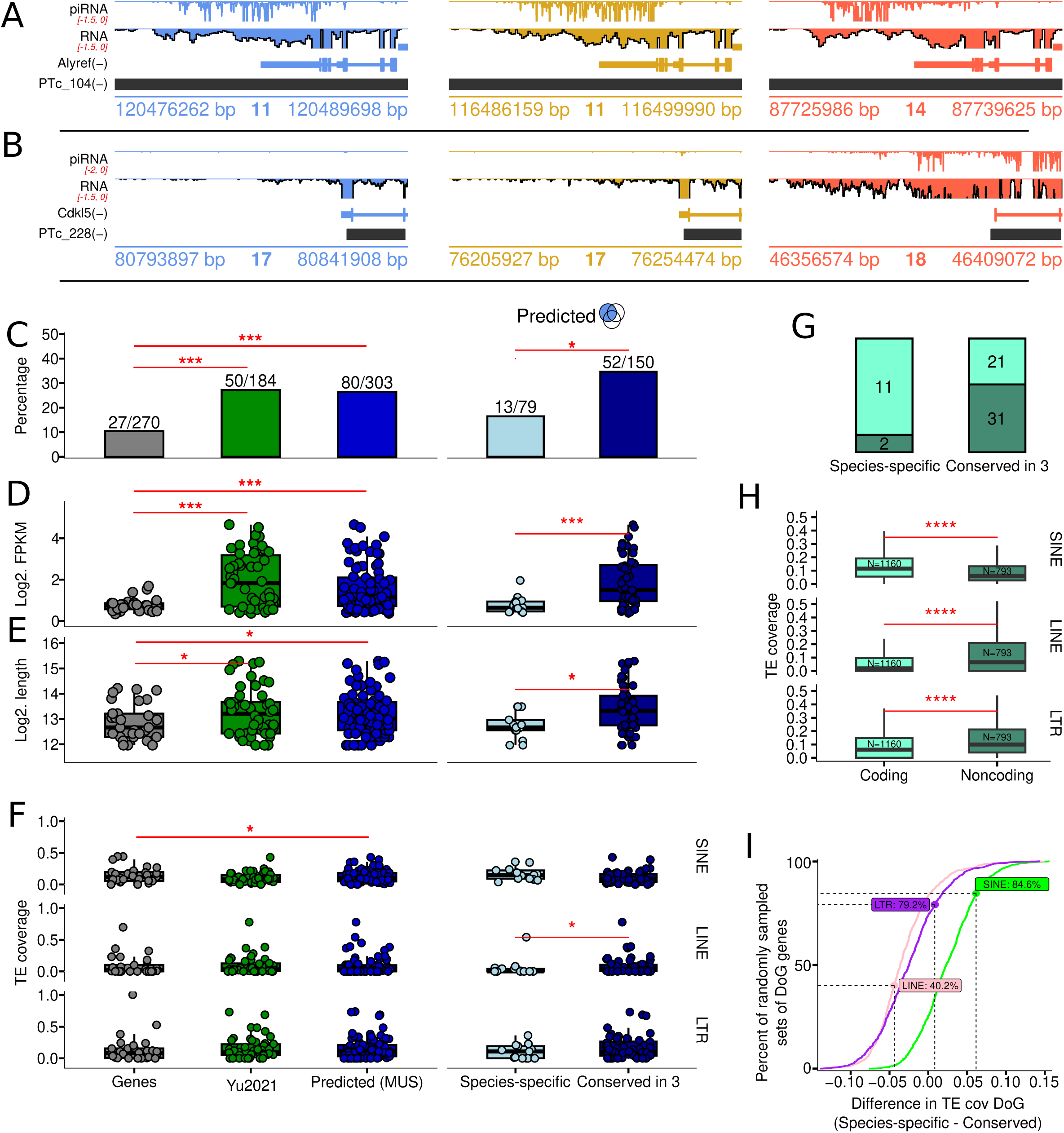
Transcriptional readthrough is more common, intense and longer in conserved compared to species-specific piRNA clusters. (A, B) Genomic tracks of conserved (A) and species-specific (B) piRNA clusters with readthrough. (C) Percentage of genes with readthrough overlapping piRNA clusters compared to expression-matched control genes with readthrough (grey data points) and Yu2021 piRNA clusters (green data points) (left), and in species-specific (light blue) versus conserved (dark blue) (right) piRNA clusters (expressed in *M. musculus*). (D) Expression (FPKM) of the downstream-of-gene (DoG) region in genes with readthrough overlapping piRNA clusters compared to control genes with readthrough (left), and in species-specific versus conserved piRNA clusters (right). (E) Length of the DoG region of genes with readthrough overlapping piRNA clusters compared to control genes (left), and in species-specific versus conserved piRNA clusters. (F) TE coverage in the DoG region of genes with readthrough overlapping piRNA clusters compared to control genes with readthrough (left), and in species-specific versus conserved piRNA clusters. (G) Proportion of coding and non-coding genes with readthrough overlapping species-specific and conserved piRNA clusters. (H) TE coverage in the DoG region of coding and non-coding genes with readthrough, regardless of piRNA production. (I) Permutation test for whether conserved and species-specific piRNA clusters differ in DoG TE coverage. For each TE class, we computed the difference in median coverage between species-specific and conserved clusters and compared it to the differences obtained from 1000 sets of random genes matched for readthrough length and coding status. Curves show the cumulative distribution of the random-set differences; the labelled percentage is the proportion of random sets with a difference smaller than the observed one. None exceeds 95%, so the differences are not significant. Positive values indicate higher coverage in species-specific clusters. Significance values in panels D, E, F and H are from Wilcoxon rank-sum (non-paired) test. Significance values in C are from Pearson’s chi-squared proportion test. Significance codes: no symbol: p ≥ 0.05. *: p<0.05, **: p<0.01, ***: p<0.001, ****: p<0.0001. Genes with readthrough, length of DoG region and expression DoG (in FPKM) were obtained from ARTDeco.

In *M. musculus*, only around 10% of expression-matched control genes have transcription DoG, whereas a significantly higher proportion (∼26%) of piRNA clusters that overlap genes have readthrough (**fig. 4C, table S9**). In addition, the RNA expression within the DoG region and the length of the readthrough region are significantly higher for genes that are piRNA clusters than for control genes (**fig. 4D-E, table S9**). Importantly, the proportion, expression and length of the readthrough region show a positive association with piRNA cluster conservation (**fig. 4C-E**).

We next examined TE coverage in DoG regions to assess the potential contribution of TEs downstream of genes to piRNA cluster conservation. Overall, we did not observe differences in DoG TE coverage (**fig. 4F, table S10**) for the major TE classes (LTRs, LINEs and SINEs) between piRNA clusters and expression-matched control genes, consistent with previous findings [29]. However, when comparing species-specific with conserved piRNA clusters, we found higher LINE coverage in DoG regions of conserved piRNA clusters compared to species-specific piRNA clusters (**fig. 4F**). Because LINEs are more common in gene-poor regions [31,32], we asked whether gene context could explain this pattern. Indeed, species-specific piRNA clusters tend to overlap more frequently with protein-coding genes with readthrough than non-coding genes with readthrough, while the opposite is true for conserved piRNA clusters (**fig. 4G**). We found that, regardless of piRNA production, DoG regions of protein-coding genes with readthrough are enriched in SINEs and depleted in LINEs and LTRs compared to non-coding genes (**fig. 4H, table S10D)**, suggesting that differences in TE content between species-specific and conserved piRNA clusters may reflect the genomic context. To test this, we asked whether the observed difference in median TE coverage of DoG regions between species-specific and conserved piRNA clusters exceeded that expected from 1000 random sets of readthrough genes with the same coding annotation and readthrough length. None of the observed differences did (**fig. 4I, table S11**). These results identify transcriptional readthrough as a feature associated with conserved piRNA clusters, but provide no evidence for a contribution of TEs downstream of genes.

### 5. piRNA expression is linked to conservation of piRNA clusters

It has been noted that some of the most highly expressed piRNA clusters are deeply conserved across mammals [10]. With a large set of conserved and species-specific clusters available to us here, we tested whether this holds more generally. Indeed, conserved piRNA clusters are among the most highly expressed ones and consistently produce more piRNAs across all three species compared to species-specific clusters (**fig. 5A-B, table S12-S13**). The same trend holds for Yu2021 piRNA clusters overlapping our set (**fig. S4A, table S14-S15**). Thus, piRNA clusters that produce abundant piRNAs tend to be conserved across species - an association we dissect below to rule out circularity arising from the use of piRNA abundance in defining piRNA clusters.

**Figure 5.**
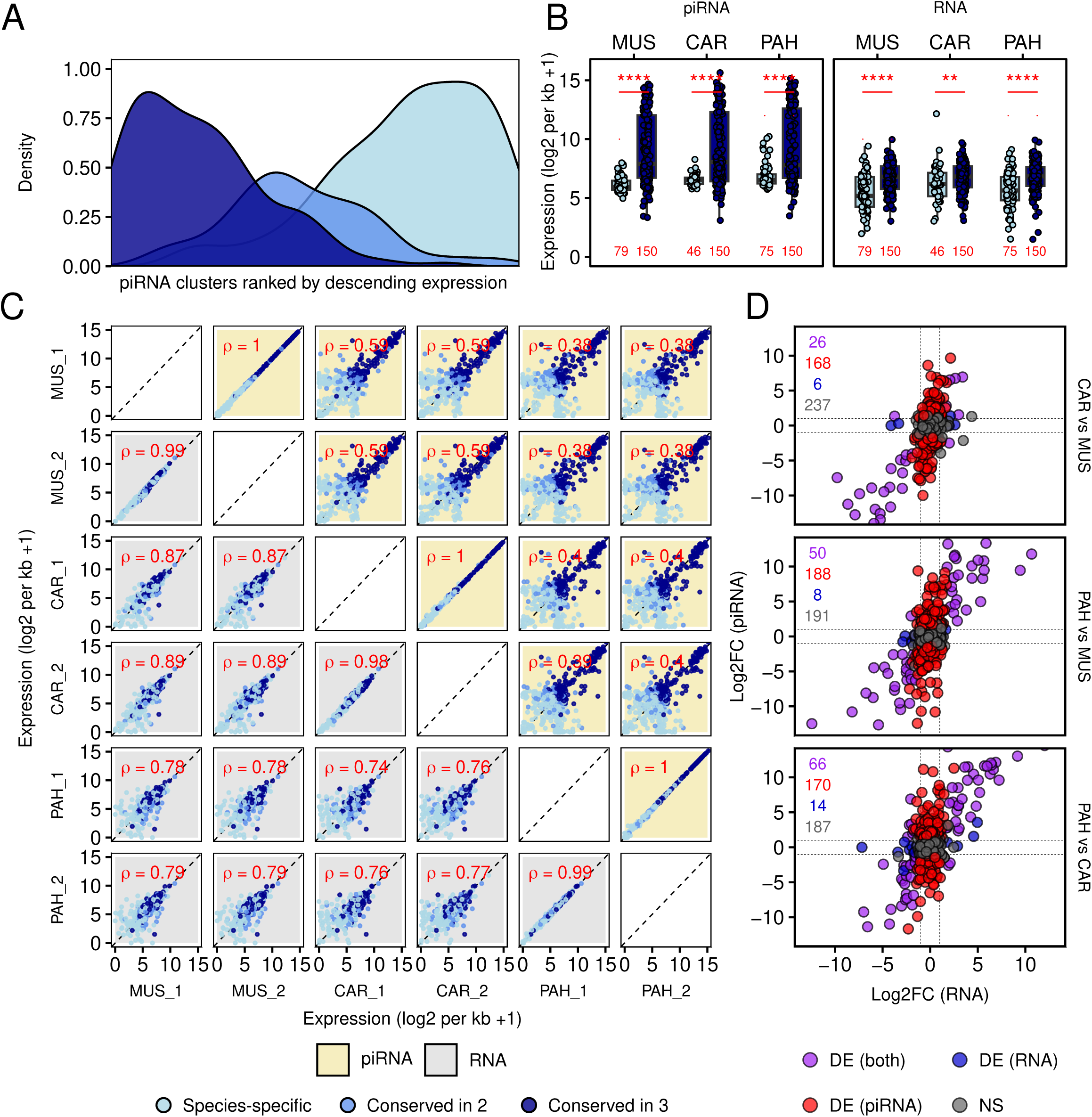
Abundance of piRNAs is linked to conservation of piRNA production. (A) Density of 437 piRNA clusters expressed in one (i.e., species-specific piRNA clusters), two or three (i.e., conserved piRNA clusters) species, ranked by descending mean piRNA expression across the three species. (B) piRNA and RNA expression of species-specific (N(MUS)=79, N(CAR)=46, N(PAH)=75) and conserved (N=150) piRNA clusters in each of the species. For species-specific piRNA clusters only those predicted in the corresponding species were taken into account. (C) Pairwise scatterplots comparing piRNA and RNA expression of 437 predicted loci between the distinct samples. Spearman correlation values are shown in red. Yellow squares are piRNA and grey are RNA. (D) Comparison of differential piRNA and RNA expression, showing that most changes between species occur at piRNA level only. Clusters were classified as differentially expressed at piRNA level, RNA level or both (absolute log2 fold change ≥ 1, adjusted p < 0.05); numbers show cluster counts per category.

Precursor expression also differs significantly between species-specific clusters and clusters conserved in all three species (**fig. 5B, table S13**), indicating that conservation of piRNA production is accompanied by conservation of precursor transcription. We therefore asked how piRNA and precursor abundance covary across species. For both, pairwise correlations are highest for the more closely related pair of species (*M. musculus - M. caroli*) and lower for the more distant pairs (*M. musculus - M. pahari* and *M. caroli - M. pahari*), as expected (**fig. 5C, fig. S4B**). However, piRNA correlations are consistently lower than the corresponding precursor correlations (piRNA Spearman’s rho 0.38-0.59 versus RNA rho 0.74-0.89) (**fig. 5C**), and decline more steeply with evolutionary distance, consistent with piRNA production being more evolutionarily labile than mRNA abundance [10].

Across all comparisons, conserved piRNA clusters show higher piRNA expression than species-specific clusters (**fig. 5B-C, fig. S4A-B**), whereas differences in precursor transcript expression are more modest (**fig. 5B-C**). Consistent with this, differential expression analysis between species reveals far more loci that change in piRNA abundance alone than in precursor expression alone or in both (**fig. 5D, fig. S4C, tables S12-S15**): across the three pairwise contrasts, piRNA-only changes outnumber precursor-only changes by 12-to 28-fold (168 versus 6, 188 versus 8, and 170 versus 14). Thus, divergence in piRNA production between species arises predominantly at the level of piRNA abundance rather than precursor transcription.

Taking the subset of our piRNA clusters overlapping with Yu2021 piRNA clusters allows us to annotate them by their timing of expression in *M. musculus* (pre-pachytene, hybrid and pachytene). Stratifying piRNA clusters this way shows that, in whole adult testis, pachytene piRNAs and their precursors are more abundant than pre-pachytene ones across all three species **(fig. S4B**), consistent with the germ cell composition of adult testis [22,33]. Across all pairwise comparisons, piRNA correlations are higher for pachytene clusters than for pre-pachytene ones, whereas the precursor correlations are higher for pre-pachytene clusters (**fig. S4B**). Thus, the greater cross-species stability of pachytene piRNA abundance is not merely a consequence of their higher abundance in adult testis, since precursor stability shows the opposite trend.

The transcription factors A-MYB and TCFL5 are transcriptional regulators of pachytene piRNA clusters in mouse [22,34], so we asked whether binding by these factors is associated with conservation. Using A-MYB and TCFL5 occupancy among the pachytene clusters annotated above [34], we found that A-MYB binds the vast majority of pachytene clusters irrespective of conservation, whereas TCFL5 binds only a minority; we found no evidence that either was associated with conservation (**fig. S4D**).

Because piRNA clusters predicted in distinct species were merged for the comparative analysis, we asked whether this step artificially inflated the length and piRNA content of conserved clusters relative to species-specific ones. Conserved piRNA clusters are indeed substantially longer than those predicted in a single species (**fig. S5A**). However, the difference in length is already present prior to merging: among the 311 *M. musculus* piRNA clusters predicted before merging (step 4 in **fig. S1E**), those overlapping conserved piRNA clusters from the final set (step 7 in **fig. S1E**) are much longer than those overlapping species-specific ones (**fig. S5B**). Because the greater length of conserved clusters is not generated by merging, their higher piRNA output is unlikely to be a mere consequence of stitching adjacent loci together. The greater length and piRNA abundance of conserved piRNA-producing loci are therefore intrinsic features rather than methodological artifacts.

To rule out the possibility that the association between piRNA expression and conservation arises from the use of piRNA expression in the definition of piRNA clusters, we tested whether the piRNA expression level of a piRNA cluster in one species (the focal species) predicts the production of piRNAs from the orthologous locus in another species (the target species). Across all six directed comparisons between species pairs, focal piRNA expression significantly predicts whether the orthologous locus is called a piRNA cluster in the target species (OR 1.49-1.97 per doubling of focal piRNA expression, all p<1e-4) (**table S16**). High piRNA expression in the focal species is therefore a robust predictor of piRNA production, and hence of cluster conservation, independently of how piRNA clusters were defined.

Since several features showed positive associations with piRNA cluster conservation, we evaluated their relative contributions using multivariate ordinal regression. For each cluster and each focal species, we classified the extent of piRNA production as an ordinal outcome: species-specific (piRNAs produced only in the focal species), close (produced in *M. musculus* and *M. caroli*), ambiguous (produced in *M. pahari* and one of the other two species, so that a gain in one lineage cannot be distinguished from a loss in the other), or conserved (piRNAs produced in all three species). Because *M. musculus* and *M. caroli* are sister species whereas *M. pahari* is the outgroup, the “close” category applies only when *M. musculus* or *M. caroli* is the focal species. Each model included piRNA expression together with additional piRNA cluster features, and the focal species as a covariate to account for the pooling of comparisons. After accounting for piRNA expression, both piRNA cluster width and precursor transcription level are significantly associated with conservation, whereas readthrough is not (**fig. 6**, **table S17**). In a model including all predictors, piRNA expression, cluster width and precursor transcription level remain significantly associated with conservation, while readthrough provides no effect beyond that of the other predictors (**fig. 6**).

**Figure 6.**
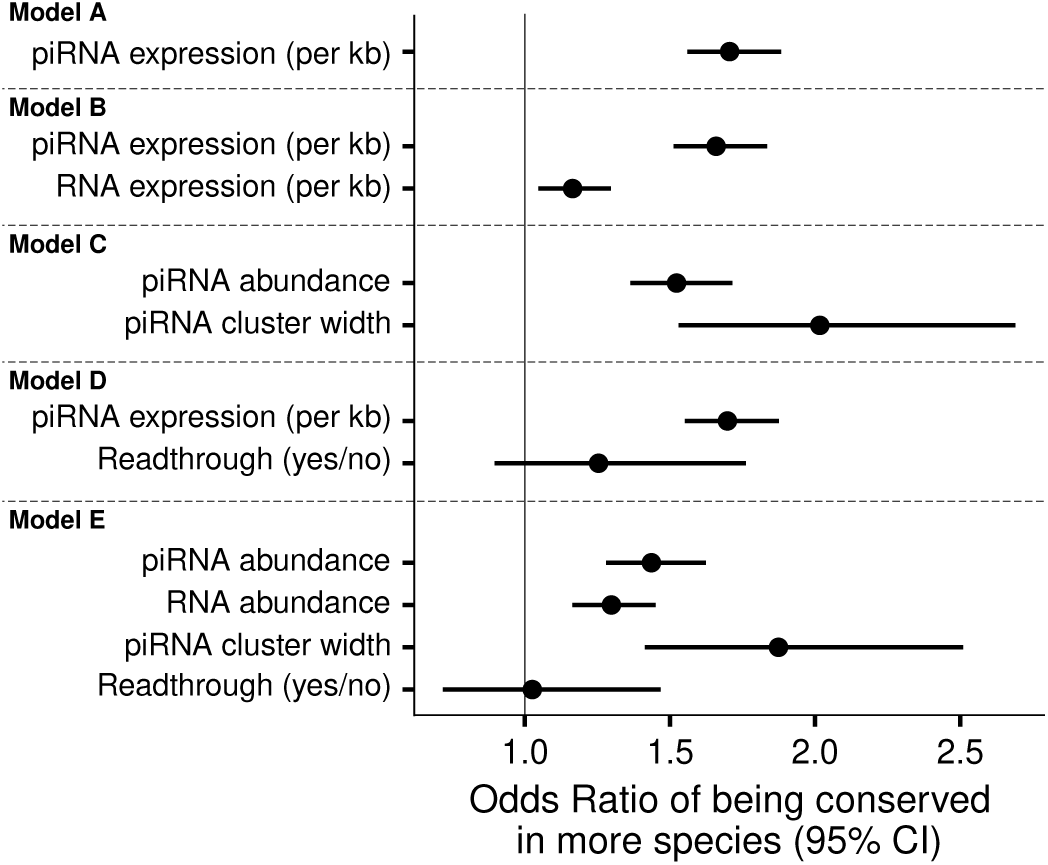
piRNA expression is the strongest predictor of piRNA cluster conservation in mouse. Odds ratios (95% confidence intervals) from ordinal logistic regression of piRNA cluster conservation level (species-specific, close, ambiguous, conserved; see Methods), per two-fold increase for continuous features. The forest plot shows piRNA expression alone, piRNA and precursor expression, piRNA expression and cluster width, piRNA expression and transcriptional readthrough and, last, all features. Expression is normalised by cluster length (per kb) except for when width is included (abundance). Models pool the three focal species and include focal species as a covariate (not shown). piRNA expression has the smallest p-value in every model. Exact values and model comparisons are in table S17.

Together, our results identify piRNA expression as the strongest predictor of piRNA cluster conservation in mouse, independently of its use in defining piRNA clusters. piRNA cluster width and precursor transcription level explain additional variation in conservation beyond that accounted for by piRNA expression, whereas readthrough does not contribute independently.

### 6. Association of piRNA expression and evolutionary persistence of piRNA clusters is maintained beyond rodents

To determine whether the association between piRNA output and cluster conservation observed in mice extends beyond rodents, we analysed human piRNA clusters previously defined by *Özata et al. (2020)* [7] (**table S18**). We stratified them according to their conservation into two classes: *human-specific* piRNA clusters, detected only in human, and *primate-conserved* clusters, detected in human, macaque and marmoset. These primate lineages diverged approximately 28 to 51 MYA [35] (**fig. 7A**).

**Figure 7.**
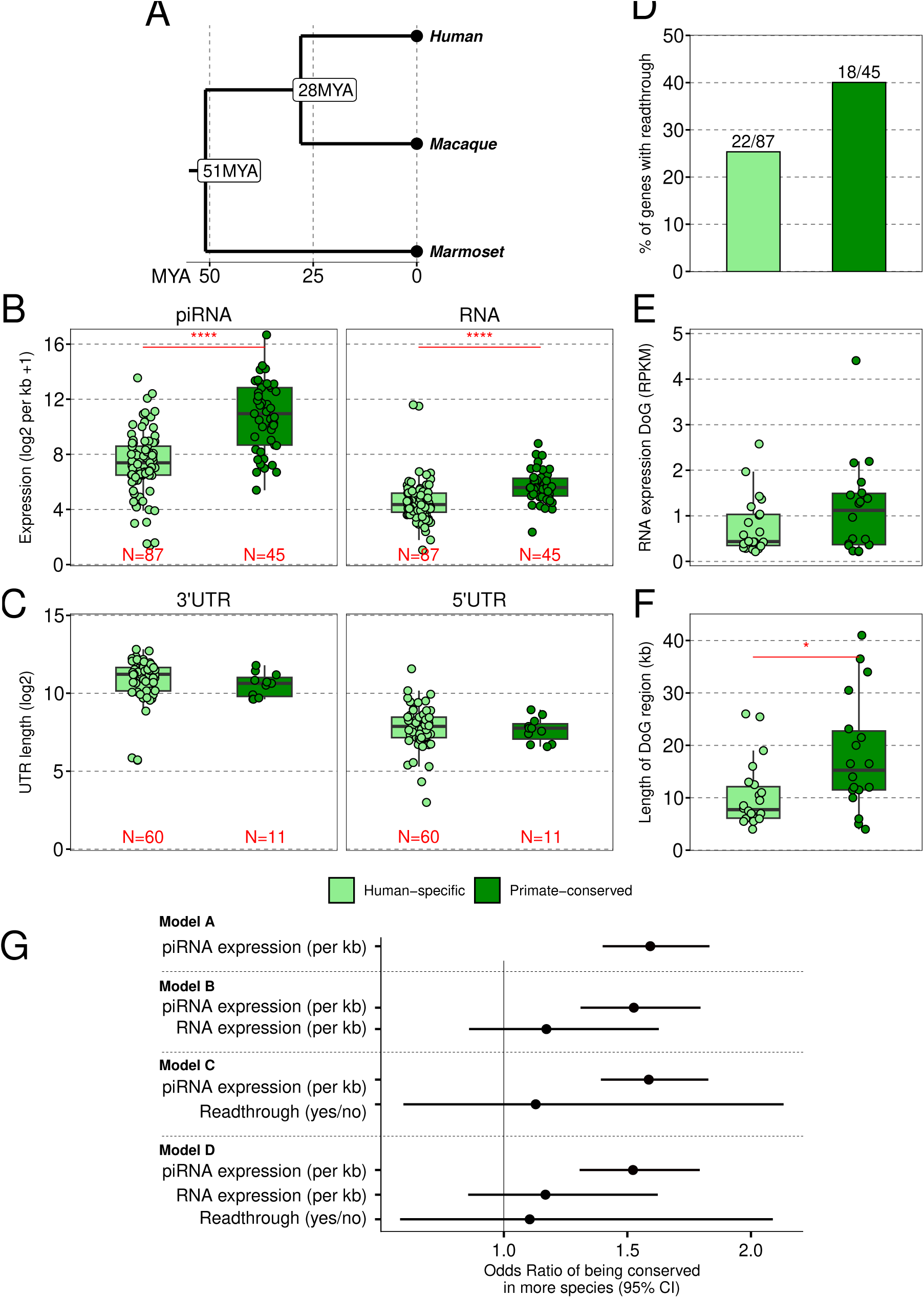
Comparison of human-specific versus primate-conserved clusters shows link between expression and conservation outside of rodents. (A) Phylogenetic tree of human, macaque and marmoset. Evolutionary distances retrieved from *Mao et al. (2024)* [35]. (B) piRNA and RNA expression of human-specific (expressed only in human, N=87) and primate-conserved (expressed in human, macaque and marmoset, N=45) piRNA clusters. (C) Length of 3’ and 5’UTRs of protein-coding human-specific (N=60) and primate-conserved (N=11) piRNA clusters. (D) Proportions of genes with readthrough overlapping human-specific (22/87) and primate-conserved (18/45) piRNA clusters. (E) Expression (FPKM) of the downstream-of-gene (DoG) region of genes with readthrough overlapping human-specific (N=22) and primate-conserved (N=18) piRNA clusters. (F) Length of the DoG region of genes with readthrough overlapping human-specific (N=22) and primate-conserved (N=18) piRNA clusters. (G) Forest plot showing odds ratios (95% confidence intervals) from ordinal logistic regression of conservation level (human-specific, close, ambiguous, primate-conserved) per two-fold increase in each feature: piRNA expression alone; piRNA expression with precursor RNA expression; piRNA expression with transcriptional readthrough; all features. Human is the only focal species. Exact values are in table S23. Significance values in panels B, E and F are from Wilcoxon rank-sum (non-paired) test and in D from Fisher’s exact test (P = 0.11). Significance codes: *: p<0.05, **: p<0.01, ***: p<0.001, ****: p<0.0001. Genes with readthrough, length of DoG region and expression DoG (in FPKM) were obtained from ARTDeco.

As in mouse, both piRNA abundance and precursor RNA abundance are higher at primate-conserved clusters than at human-specific loci (**fig. 7B, table S19, S20**). Consistent with mouse, neither 3′UTR nor 5′UTR length differs significantly between the two classes (**fig. 7C, table S21**). We next examined transcriptional readthrough in piRNA-producing genes. The proportion of piRNA clusters with detectable DoG transcription and the level of transcriptional readthrough do not differ significantly between classes (**fig. 7D-E, table S22**). Only the length of the DoG region is significantly greater in primate-conserved clusters (**fig. 7F**).

To evaluate the relative contributions of piRNA abundance, precursor abundance and readthrough in conservation, we fitted ordinal logistic regressions to model piRNA cluster conservation based on each of these features (**fig. 7G, table S23**). We split the piRNA clusters in human-specific, close (expressed in human and macaque), ambiguous (expressed in human and marmoset), and primate-conserved (expressed in all three primates). Across all models tested, piRNA abundance is a significant predictor of conservation, whereas neither precursor RNA abundance nor the presence of transcriptional readthrough reaches statistical significance when considered alongside piRNA abundance (**fig. 7G**.

Despite representing a larger evolutionary distance than that separating our three mouse species, these results extend our findings in mouse and support a conserved link between high piRNA expression and the evolutionary persistence of piRNA clusters in primates as well as rodents, suggesting that this relationship may hold broadly across mammals.

## DISCUSSION

In this study, we show that evolutionarily persistent piRNA-producing loci are those that generate the highest levels of piRNAs. Among all the genomic features tested, piRNA abundance was the strongest predictor of conservation. This relationship was observed independently in both rodents and primates, suggesting that piRNA output is a general predictor of conservation of piRNA production across mammalian evolution.

Our study further highlights that, out of the hundreds of loci producing piRNAs in a given species, only a small subset is protected from evolutionary drift. The rapid loss of the remaining loci suggests that either the fitness benefit they confer is transient or, more simply, that their piRNA output does not provide a significant selective advantage. Efficient piRNA-mediated silencing depends on piRNA abundance relative to the target, in flies [36] and in mouse spermatocytes [37,38]. Only a small fraction of piRNAs reach high abundance [37–39]. In agreement with these observations, we find that loci producing highly abundant piRNAs are preferentially conserved. Interestingly, this parallels a well-established observation in protein-coding gene evolution: highly expressed genes are less likely to be lost across species and evolve more slowly than lowly expressed ones [40,41]. Although protein-coding genes and piRNA-producing loci are fundamentally different genomic entities, the same relationship between expression level and evolutionary persistence appears to hold for both. This could reflect that higher expression amplifies functional impact through increased molecular interactions, making the locus more likely to be under strong purifying selection. In the case of piRNAs, a higher piRNA output increases the probability that a piRNA cluster produces a functional piRNA at the abundance required for target silencing. Similarly, a wider piRNA-producing region, which we found to significantly explain conservation, would increase the diversity of piRNA sequences, and a higher transcription level would increase the abundance of individual piRNAs. In addition, a minimal expression level, likely set by the abundance of the target, may be required for a cluster to be functional, after which selection could favour variants that further increase its expression, so that high output would also be, in part, a consequence of persistence.

The strong association between conservation and piRNA abundance argues against clearance of the precursor transcripts themselves as the reason why high-output loci are conserved, since the efficacy of such clearing would be largely independent of how many piRNAs a locus produces. Moreover, piRNA production carries a cost (precursor transcription and processing) that scales with output. Such a cost could be sufficient to select against high-output loci that serve no purpose, but too small to purge the low-output majority. Collectively, these observations suggest that the majority of piRNAs expressed in mammalian testis likely represent tolerated molecular noise rather than functionally important regulatory molecules, while the highly expressed, conserved minority are those most likely to be functional.

Recently, transcriptional readthrough has been found to be associated with piRNA production downstream of genes [29]. Testis in general is a tissue with higher frequency of readthrough transcription compared to other tissues [42]. We found that species-specific piRNA clusters and expression-matched control genes show similar levels of readthrough, while conserved piRNA clusters have higher transcriptional readthrough. It has been proposed that piRNA production from some readthrough transcripts can silence transposons and viral elements inserted downstream of genes [29,43]. After accounting for gene type and readthrough length, we did not find a significant enrichment of any of the major TE classes in or downstream of species-specific piRNA clusters compared to conserved ones. Importantly, once we accounted for piRNA expression and other piRNA cluster features, readthrough did not explain conservation of piRNA production in regression models. This suggests that readthrough is a byproduct of something that underlies the elevated output of conserved loci, rather than an independent contributor to their persistence. Consistent with this, piRNA clusters with transcriptional readthrough commonly initiate piRNA production within the gene (**fig. 4A-B**) and extend it further downstream. Considering that pachytene piRNA precursors tend to have atypically long and often unspliced transcripts [23]and that inefficient splicing is linked to transcriptional readthrough [44], perhaps transcriptional readthrough is a consequence of the transcriptional features of piRNA precursors, rather than directly linked to piRNA production.

One limitation of this study is that it is based on small RNAs from whole testis of adult mice, which are enriched in pachytene cells; it therefore remains unclear whether piRNA expression also explains the conservation of pre-pachytene piRNA clusters. Second, although we tested the features most plausibly linked to piRNA function in our analyses, piRNA abundance remained the strongest predictor of conservation. That the dominant signal is the quantity of piRNAs produced, rather than a sequence feature, is itself informative, though it does not formally resolve whether selection acts on piRNA output directly or on the process sustaining high production. Finally, because piRNA-producing loci vary between strains of the same species [18], and this variation is predominantly strain-specific gains, some piRNA clusters scored as species-specific may reflect strain-specific piRNA production rather than true species differences, since our calls were made on a single representative strain per species. There is no a priori expectation that such loci differ in expression from the rest, so this is unlikely to bias the expression-conservation relationship in a particular direction.

## CONCLUSIONS

In conclusion, by analysing small RNAs from testis of closely related mouse species, and extending our analysis to primates, we identified features related to the conservation of piRNA production. The greater conservation of piRNA clusters with higher piRNA output may be due to critical functional roles, either through more piRNA-target interactions or through more robust silencing of their targets. Overall, our findings reveal that the abundance of piRNAs produced from a locus is the strongest predictor of conservation of piRNA production in mammals.

## MATERIALS AND METHODS

### RNA extraction, library construction and sequencing

RNA was extracted from whole testes from *Mus musculus* strain C57BL/6J, *Mus caroli* strain CAROLI/EiJ and *Mus pahari* strain PAHARI/EiJ. RNA was extracted using RNA TRIzol^TM^ PLUS Reagent (Thermo Fisher Scientific, ref 12183555). Small RNA libraries were prepared following the TruSeq^TM^ protocol with automated pooled library size selection using Pippin prep. All small RNA-seq samples were sequenced together on one lane on HiSeq2500 from Illumina (1×50nt sequencing mode). Total RNA sequencing libraries were prepared with the TruSeq Stranded Total RNA Library Prep Gold Kit and sequenced on NovaSeq6000 (2×150nt sequencing mode).

### Genome assemblies, gene and repeat annotations

Genome assemblies and gene annotations were downloaded from Ensembl, release 104 (Dyer2024): GRCm39/mm39 for *Mus musculus,* CAROLI_EiJ_v1.1 for *Mus caroli and* PAHARI_EiJ_v1.1 for *Mus pahari.* RepeatMasker annotations were retrieved from UCSC Genome Browser [45] corresponding to assemblies GRCm39/GCF_000001635.27 for *Mus musculus,* CAROLI_EiJ_v1.1/GCF_900094665.1 for *Mus caroli and* PAHARI_EiJ_v1.1/GCF_900095145.1 for *Mus pahari*. The gold standard set of mouse piRNA clusters (strain C57BL/6J) was retrieved from *Yu et al. (2021)* [23].

### RNA-seq and small-RNA seq basic data processing

RNA-seq reads were mapped to the corresponding assembly using STAR [46]. PCR duplicates were removed with samblaster v0.1.26 [47]. For the small RNA-seq data, the 3’ adapter (TGGAATTCTCGGGTGCCAAGG) was trimmed using *Cutadapt* v3.7 [48], requiring a 9nt match with the adapter sequence. Reads shorter than 19nt or longer than 36nt and any reads not matching the adapter were discarded. Then, reads were filtered based on quality using the FASTX Toolkit v0.0.14 (http://hannonlab.cshl.edu/fastx_toolkit/) allowing minimum quality score 30 over at least 90% of nucleotides. Trimmed and filtered reads were mapped to the corresponding assembly for each species using *Bowtie* v1.3.1 [49], retaining uniquely mapping reads and allowing one mismatch. For single-end small RNA-seq, reads were separated by strand and coverage tracks were generated with *bamCoverage* (*deepTools* [50]), normalised to counts per million (CPM) at 1 bp resolution. For paired-end RNA-seq data, strand-specific CPM-normalised coverage was generated at 100 bp resolution.

### Identification of orthologous regions

Because data from different species were aligned to different genome assemblies, coordinates were converted between assemblies. For conversions within *Mus musculus* species (e.g., GRCm38/mm10 to GRCm39/mm39), we used liftOver [51]. For conversions between species (e.g., GRCm39/mm39 to CAROLI_EiJ_v1.1), we used the ENSEMBL Compara Perl API [52], from which we obtained the genomic coordinates and the multiple sequence alignment of the query region in the desired species.

### Prediction of piRNA-producing loci

We used the proTRAC pipeline [53] to predict piRNA producing loci in each sample of the three mouse species based on the small RNA-seq data. To process the reads we used the *NGS toolbox* v2.1 [54] to collapse (*TBr2_collapse.pl*) and remove low-complexity (*TBr2_duster.pl*) small RNA-seq reads. Processed reads were mapped to the corresponding assembly for each species using *sRNAmapper* v1.0.4 and multi-mapping reads were weighted based on the uniquely-mapping reads in the region using *reallocate* v1.1. Finally, we predicted piRNA clusters using proTRAC v2.4.4 [53]with default options. The bidirectional clusters were split using the coordinates given by proTRAC, and those clusters covered by repeats by more than 80% were removed. To create a common piRNA cluster annotation for the three species, we converted the predicted clusters from each corresponding genome to GRCm39/mm39 using the ENSEMBL Compara Perl API. Then, piRNA clusters from the same species were merged with *BEDTools* v2.30.0 [55,56], forcing the same strandedness and a maximum distance of 10kb, and only those clusters present in both replicates within the same species were retained, leading to 311, 216 and 246 predicted piRNA clusters in *M. musculus*, *M. caroli* and *M. pahari*, respectively. Finally, we merged the three piRNA cluster sets from the distinct species being within 10kb of each other and on the same strand, resulting in 513 piRNA clusters that were converted back to the distinct assemblies using ENSEMBL Compara Perl API. These loci were named according to the order of appearance in the *M. musculus* genome (sorted by chromosome and start coordinate). Only those loci with one-to-one orthologs between the three assemblies were retained for further analyses, which led to a set of 471 piRNA clusters. These 471 loci were predicted by proTRAC in one, two or three species. Coordinate transfer between assemblies and merging of per-species clusters changed the length of the final loci and their read counts. We therefore applied a second-pass filter to refine the set of piRNA-producing loci in each species: ranking loci by ascending piRNA expression against the proTRAC calls revealed a clear inflexion point in each species, which we used as a per-species expression threshold to define whether a locus produces piRNAs (**fig. S6**). This resulted in 437 loci defined as piRNA-producing in one, two or three species.

### Quality control of piRNA-like traits and piRNA cluster prediction

We computed the length, first and tenth nucleotides, and first codon composition of trimmed and quality filtered small RNA-seq reads using custom AWK commands. We counted the 5’-to-5’ overlapping distance between reads in opposite strands with *TBr2_pingpong.pl* from the *NGS Toolbox* v2.1 [54], using the mapped reads used for the proTRAC pipeline. To assess the quality of the predicted piRNA clusters, we compared the 311 piRNA clusters predicted in the two samples of *Mus musculus* (fig. S1E) and the 215 Yu2021 piRNA clusters. Small RNA-seq reads mapping in the clusters were counted using *featureCounts*, requiring a minimum mapping quality of 1 and a minimum overlap of 18nt with the cluster, allowing reads to overlap more than one feature, and ignoring strand. Proportions of coding and bidirectional (i.e., divergently transcribed) piRNA clusters were computed as in the corresponding methods sections. The 311 predicted piRNA clusters were intersected to the 215 Yu2021 piRNA clusters. The predicted piRNA clusters not overlapping to Yu2021 set were intersected to 900 piRNA clusters previously predicted using distinct mouse strains and tissues [18]. The class of the predicted piRNA clusters in terms of the timing of expression (i.e., pre-pachytene, hybrid, pachytene) was defined based on the overlap to Yu2021 piRNA clusters.

### Estimation of expression and differential expression analysis of piRNA clusters

To quantify the expression of piRNA clusters, we counted reads mapping to the piRNA clusters using *featureCounts* v2.0.1 [57]. For small RNA-seq, we required a minimum alignment quality of 1 and a minimum 18nt overlap with the feature and allowed multiple overlaps. For long RNA-seq, we used the same requirements as in small RNA-seq, and included the flag for paired-end data, requiring both ends to be mapped and counting fragments instead of reads. For normalization and differential expression analysis, we used *DESeq2* v1.38.3 [58], requiring an absolute log_2_(fold-change) value of 1 and a false discovery rate (FDR or adjusted p-value) of 0.05 to define a piRNA cluster as differentially expressed (DE). To account for differences in length between piRNA clusters and species, RNA and piRNA expression values were calculated dividing the DESeq2-normalized counts by the length of the piRNA clusters in kilobases (hereafter expression per kb; expression not divided by length is referred to as abundance).

### Annotation of genic and bidirectional piRNA clusters

piRNA-producing loci can be protein-coding genes, annotated non-coding genes and regions not overlapping annotated genes. To annotate piRNA clusters according to the overlap with genes, we used bedtools to intersect the coordinates of the 471 loci (before the second pass filter) to protein-coding and long non-coding RNA (lncRNA) transcripts and 3’UTRs in the *M. musculus* genome. First, we annotated 3’UTR loci by intersecting them to 3’UTRs of coding genes, requiring the same strandedness and a reciprocal overlap fraction between predicted loci and 3’UTRs of 0.33. If a locus overlapped more than one gene, we retained the hit with the maximum overlap among all. For the rest of the coding loci, we intersected them to protein-coding transcripts using the same approach as in 3’UTR piRNA clusters. Since some predicted genic loci are small and do not fulfil an overlap fraction of 0.33, we repeated the intersection by allowing a fraction of piRNA cluster overlap of 0.9 and a fraction of gene overlap of 0.1, so they were correctly annotated as coding. Next, we annotated non-coding loci by intersecting them to lncRNA transcripts using the same approach as for the coding piRNA clusters. Finally, we labelled the loci not overlapping protein-coding or lncRNA transcripts as intergenic and found the closest upstream and downstream genes using *bedtools*.

To annotate bidirectional piRNA clusters, we used a custom R function that counts the distance between consecutive piRNA clusters and assigns them as bidirectional if they are separated by a distance shorter than 500bp.

### TE overlap to piRNA clusters and permutation test

To compute the coverage of TEs in piRNA clusters and genes we used *bedtools intersect* previously extending predicted piRNA clusters by 5kb upstream and downstream. For each transposon class (i.e., SINEs, LINEs, LTRs), we computed the coverage (i.e., bases of overlap divided by the length of the piRNA cluster or gene) in each gene or piRNA cluster. To test whether individual TE families were enriched in sets of predicted piRNA clusters, we performed a permutation test (1000 permutations) with the *crosswisePermTest* function from the *regioneReloaded* R package, using *circularRandomizeRegions* as the randomisation function and numOverlaps as the evaluation function [59]. The test was restricted to individual

TE families (within SINEs, LINEs and LTRs) with more than four insertions across the full piRNA cluster set. The adjustment method for p-values was the Benjamini-Hochberg algorithm.

### piRNA cluster expression-matched control genes

To compare genic piRNA clusters against other genes, we took a subset of random non piRNA-producing genes with similar expression distribution to genes overlapping annotated piRNA clusters (**table S24**). Briefly, we estimated expression of all genes in our RNA-seq data from *Mus musculus* using *featureCounts*, requiring a minimum alignment quality of 1, both ends to be mapped. We counted fragments mapping to transcripts and summarized them by gene. We filtered genes to get only protein-coding and long non-coding RNA (lncRNA) genes, and we calculated the mean FPKM value of both *Mus musculus* samples for each gene. Then, genes overlapping the 215 Yu2021 and 437 predicted piRNA clusters (referred to as reference set; 270 genes total, see “Annotation of genic and bidirectional piRNA clusters”, above) were separated from those not overlapping any piRNA clusters (referred to as control gene set). The expression range of control genes was constrained to the minimum and maximum values observed in the reference set. Expression values of the reference set of genes were divided into 100 bins, which were then used to binarize the control gene expression data. We counted the number of genes per bin and randomly resampled control genes within each bin to match the reference set expression distribution (**fig. S7**).

### UTR length, AU-rich element and TE enrichment in UTRs

To get the UTR length of genic piRNA clusters and other genes, we retrieved the length of the UTRs of each transcript. For each gene, we took the mean length of all its transcripts. To study the AU-rich element (ARE) enrichment in UTRs of genic piRNA clusters and other genes, for each species we downloaded the UTR sequences of all transcripts from ENSEMBL (release 104) using *biomaRt* R package [60]. We calculated the AREscore [61] for each transcript using a custom reimplementation of the *hpgl_arescore()* function from the R package *hpgltools* (https://github.com/elsayed-lab/hpgltools/) and the *AREscore()* R function from *SeqTools* (https://github.com/lianos/seqtools/). For each gene, we took the mean AREscore of all its transcripts. To assess the TE enrichment in UTRs, we used *bedtools intersect* to overlap UTRs to transposons from repeatMasker annotation. For each transposon class, we computed the coverage in each transcript. For each gene, we took the mean TE coverage of all its transcripts. When using data from only one species, predicted piRNA clusters were filtered to show only piRNA clusters originally predicted at least in that species. Only genes with both annotated 5’ and 3’UTRs that could be retrieved were used for this analysis. To compare genes across species, only UTRs of genes with one-to-one orthologs across *M. musculus*, *M. caroli* and *M. pahari* were used.

### Transcriptional readthrough and TE enrichment downstream of gene

To analyse genes with transcriptional readthrough, we used ARTDeco v0.4 [62] pipeline. Total RNA-seq from all samples was used as input for ARTDeco, using gene annotation and chromosome sizes from the corresponding species, requesting a readthrough distance of 5000 bp and a read-in distance (i.e., distance to consider a gene to be transcribed from the upstream gene) of 1000bp. To define piRNA clusters with readthrough, we intersected the coordinates of predicted piRNA clusters to those of genes with readthrough - including the downstream-of-gene (DoG) region-as defined by ARTDeco. For control genes with matched gene expression to piRNA clusters, we directly took the genes with readthrough given by ARTDeco. Readthrough length and expression (FPKM) were taken from ARTDeco. To compute the TE coverage, SINEs, LINEs and LTR repeats from repeatMasker were overlapped to the given readthrough regions using *bedtools intersect*.

To assess whether the observed differences in TE enrichment between species-specific and conserved piRNA clusters could be explained by gene type (coding or non-coding) or readthrough length, we performed a permutation-based randomization test with 1000 permutations comparing the 80 predicted piRNA clusters with readthrough in *M. musculus* with matched random regions. From the 1860 genes with readthrough not overlapping the predicted piRNA clusters (1111 coding, 749 noncoding), we randomly sampled 1000 sets of readthrough genes with the same number of coding and non-coding genes as the real set (40 coding, 40 noncoding) and assigned the readthrough length and conservation status of each real gene to the corresponding sampled gene. The TE coverage was calculated for each of the readthrough regions, and the median TE coverage of species-specific and conserved piRNA clusters was taken for the observed and randomised datasets in each permutation. We calculated the difference of medians between species-specific and conserved piRNA clusters and computed the proportion of randomised sets whose difference of medians was lower or higher than that of the observed set. If the percentage of lower or higher sets was over 95%, we concluded that the difference in TE coverage between species-specific and conserved piRNA clusters was significant.

### Analysis of A-MYB and TCFL5-bound Yu2021 piRNA clusters

To define which mouse pachytene piRNA clusters were bound by A-MYB and TCFL5, we retrieved the distances from the piRNA clusters TSS to the nearest ChIP-seq peak summit from *Yu et al. (2023)* [34]. Those piRNA clusters whose TSS was closer than 500bp to the nearest summit were considered bound by the corresponding transcription factor.

### Analysis of human piRNA clusters

We downloaded 182 human piRNA clusters previously defined by *Özata et al*. *(2020)* [7] in GRCh37/hg19 coordinate system. To stratify them by conservation level, we took the human piRNA clusters expressed in macaque and marmoset from *Özata et al. (2020)* [7] Human piRNA clusters expressed in human, macaque and marmoset were defined as primate-conserved, while those only expressed in human were defined as human-specific.

Human bulk RNA-seq (n=14, GSM4030196-GSM4030209) and small RNA-seq (n=13, GSM4030214-GSM4030218, GSM4030220-GSM4030227) data generated by *Özata et al. (2020)* [7] from adult human testis were downloaded from GEO (GSE135791) and were analysed the same way as the rest of the sequencing data, with small variations. The 3’ adapter for small RNA-seq data was NNNTGGAATTCTCGGGTGCCAAGG, the pre-processed reads were mapped to GRCh38/hg38 human genome assembly (ENSEMBL release 104), and GRCh37/hg19 piRNA clusters were converted to GRCh38/hg38 coordinate system for downstream analysis using liftOver. Transcriptional readthrough in human piRNA-producing genes was analysed with ARTDeco as described for mouse, using the human RNA-seq data and GRCh38/hg38 gene annotation.

### Logistic regression models

To assess the contribution of distinct features of piRNA clusters in one species (referred to as focal species) to its expression in the other species (referred to as target species) or the conservation level of the piRNA cluster, we built different logistic regression models. The variables in the model were transformed to have a similar range of values. RNA and piRNA expression (DESeq2-normalized counts), either normalized by length (e.g., piRNA expression (per kb)) or not (e.g., piRNA abundance), were transformed by a base-2 logarithm, and so was done for piRNA cluster width. For transcriptional readthrough, we considered a binary variable indicating whether or not the piRNA cluster overlaps a gene with readthrough, since including both expression and length of the DoG region led to collinearity between these two variables.

Six directed pairwise binomial logistic regression models were built using the R function *glm(family=”binomial”)* to assess whether piRNA expression (i.e., independent variable) in one species predicted the presence of the piRNA cluster (i.e., response variable) in the other species. Using the same focal-target pairwise data, we also computed the Spearman’s correlation of piRNA cluster expression in focal and the piRNA cluster expression in the target.

To include the distinct features and predict conservation levels, we used ordinal logistic regression models using the *clm()* function from the *ordinal* R package [63]. For each focal species, we classified piRNA clusters in each focal species into species-specific (expressed only in the focal species), close (expressed in *M. musculus* and *M. caroli*), ambiguous (expressed in *M. pahari* and one of the other species) or conserved (expressed in the three species).

For human piRNA cluster conservation, we built ordinal logistic regression models as in mouse, but using human as the only focal species. We classified piRNA clusters in human-specific (expressed only in human), close (expressed in human and macaque), ambiguous (expressed in human and marmoset) and primate-conserved (expressed in the three species).

## Availability of data and materials

All small RNA-seq and RNA-seq samples are deposited to Gene Expression Omnibus (GEO, GSE344279). Custom scripts written to analyze the data are available from https://github.com/amitjavilaventura/Mitjavila_et_al_2026_rodent_pirna

## Authors’ contributions

Conceptualization: Adrià Mitjavila-Ventura (AMV) and Tanya Vavouri (TV); Data curation: AMV and TV; Formal analysis: AMV and TV; Funding acquisition: AMV and TV; Investigation: AMV and TV; Methodology: AMV and TV; Resources: AMV, Samuele M. Amante (SMA), Cristina Moreta-Moraleda (CMM) and TV; Writing – original draft: AMV and TV; Project administration: TV; Supervision: TV, Miguel R. Branco (MRB) and Sonia V. Forcales (SVF).

## Use of generative AI and AI-assisted technologies

During the preparation of this manuscript, the authors used ChatGPT (OpenAI) and Claude (Anthropic) to assist with language editing and clarity of the text. Reviewer3 (https://reviewer3.com/) was used to identify critical gaps in an early version of the manuscript that we subsequently addressed. These tools were not used to generate original scientific content or interpretations. After using these tools, the authors reviewed and edited the content as needed and take full responsibility for the content of the publication.

## Acknowledgements

We thank the IGTP High Content Genomics and Bioinformatics Unit and their staff for their contribution and technical support.

## SUPPLEMENTARY FIGURES

**Supplementary figure 1.**
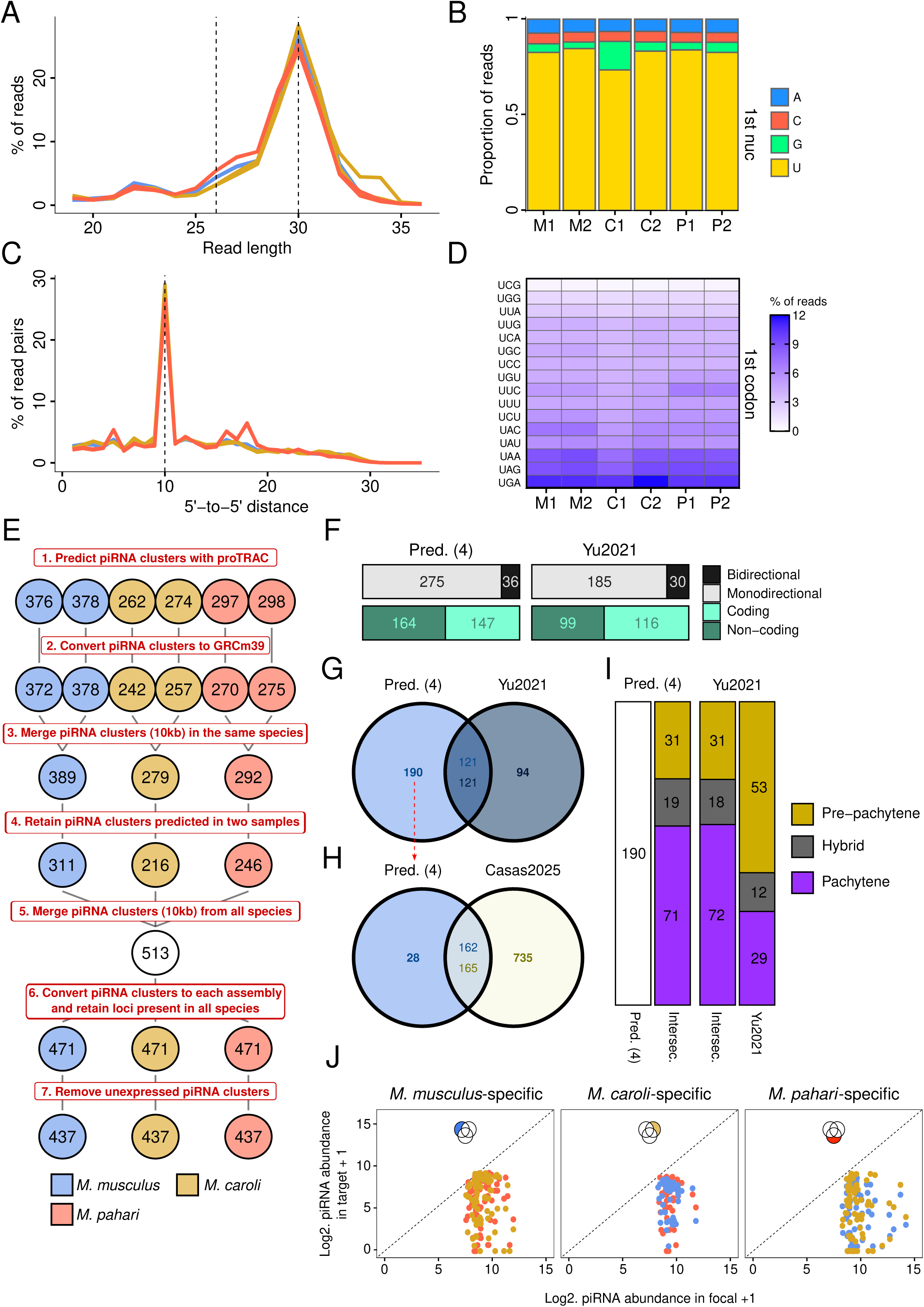
Quality assessment of small RNA-seq reads and prediction of piRNA clusters. (A) Read length distribution of reads in small RNA-seq libraries. Vertical dashed lines show 26 and 30 nt. (B) 1st nucleotide composition of reads in small RNA-seq libraries. (C) 5’-to-5’ distance of antisense overlapping reads in small RNA-seq libraries. (D) Proportion of U-starting codons in the first three nucleotides of reads in small RNA-seq libraries. (E) Pipeline for piRNA cluster prediction. Numbers after each step are indicated. For panels F-I, the 311 piRNA clusters predicted in M. musculus (step 4, blue) were used. (F) Proportion of monodirectional versus bidirectional, and coding versus non-coding piRNA clusters predicted by us and defined in M. musculus by Yu2021 (referred to as Yu2021 piRNA clusters). Directionality and coding status were defined using custom functions. (G) Overlap of 311 predicted piRNA clusters and 215 Yu2021 piRNA clusters. (H) Overlap of predicted piRNA clusters not overlapping Yu2021 piRNA clusters and piRNA clusters defined using a set of 5 inbred mouse strains by *Casas et al. (2025)* (referred to as Casas2025 piRNA clusters). Note that in this dataset, piRNA clusters must be predicted in two samples from each species to be retained, while in Casas2025 piRNA clusters, all piRNA clusters from all samples were retained in the final piRNA cluster set. (I) Class of predicted and Yu2021 piRNA clusters. For predicted piRNA clusters, the class of the overlapping Yu2021 piRNA cluster was used. (J) piRNA expression of the final piRNA clusters predicted in only one species, showing that abundance of piRNAs in the predicted species is higher compared to the others.

**Supplementary figure 2.**
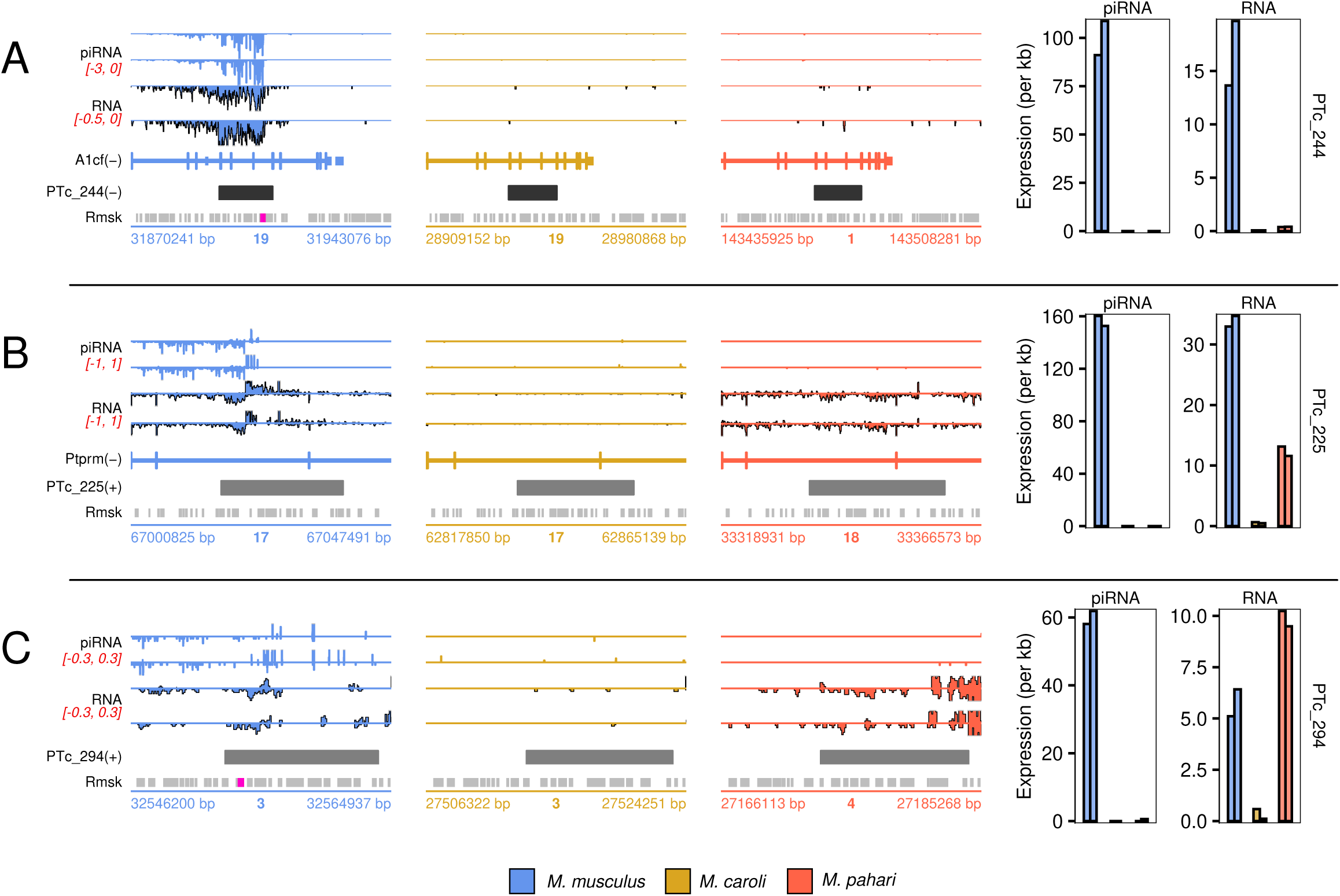
Polymorphic TEs may drive polymorphic piRNA production across the three mouse species. Genomic tracks (left) and piRNA and RNA expression (right; *M. musculus*, *M. caroli* and *M. pahari*, in this order) of three piRNA clusters with polymorphic TE insertions across the three mouse species: (A) PTc_244, sense to A1cf; (B) PTc_225, antisense to Ptprm; (C) PTc_294, an intergenic locus on chromosome 3 between *Kcnmb3* and *Zfp639*. Barplots show individual replicates. TE insertions likely associated with piRNA production are highlighted in purple. Repeats were retrieved from the RepeatMasker track of the corresponding species.

**Supplementary figure 3.**
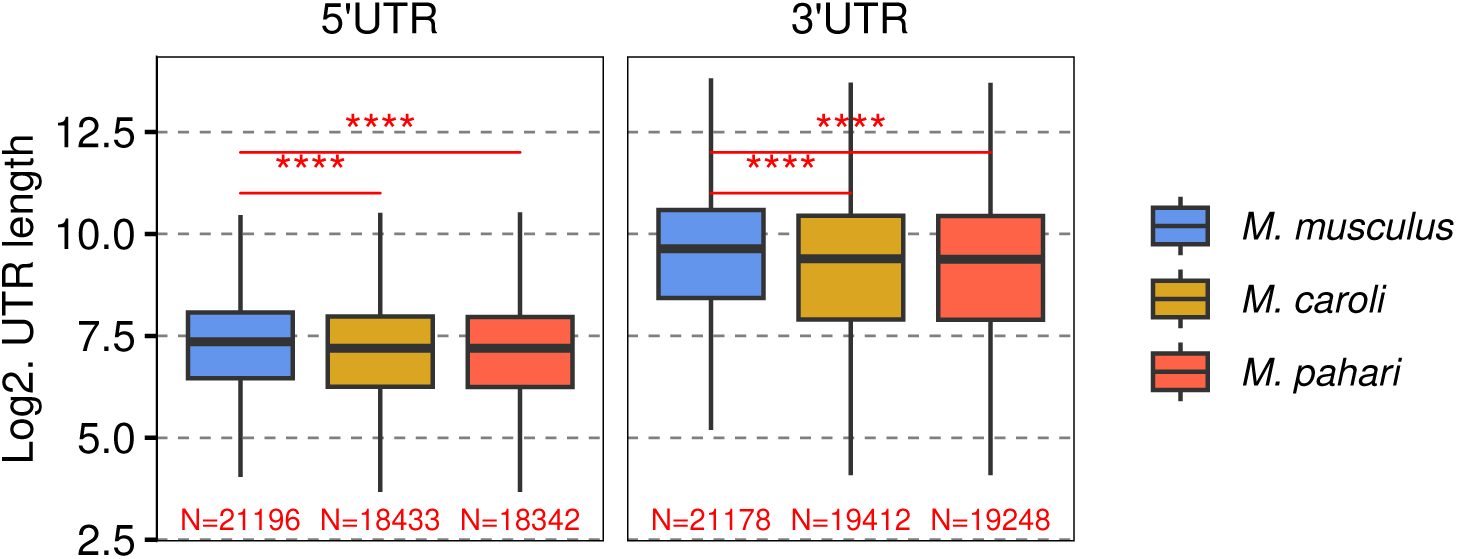
UTR length of all genes in *M. musculus, M. caroli* and *M. pahari*shows generally longer UTRs in *M. musculus*. Significance values are from Wilcoxon rank sum (non-paired) test. Significance codes: *: p<0.05, **: p<0.01, ***: p<0.001, ****: p<0.0001.

**Supplementary figure 4.**
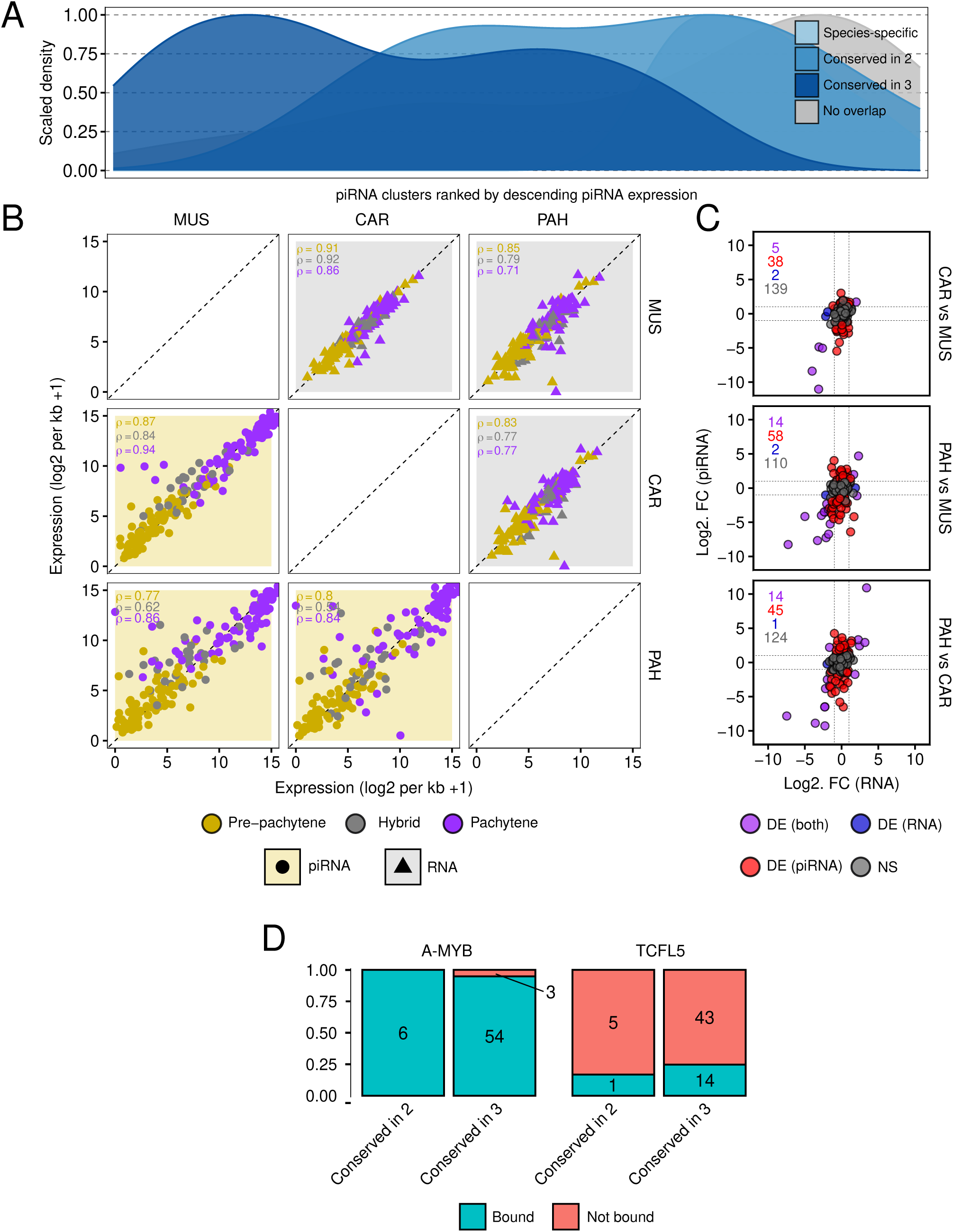
piRNA and RNA expression in Yu2021 piRNA clusters, and their binding by A-MYB and TCFL5. (A) Scaled density of 184 Yu2021 piRNA clusters that are species-specific or conserved in two or three mouse species based on overlap with 437 predicted piRNA clusters, ranked by descending mean expression across the 3 species. (B) Pairwise scatterplot comparing the piRNA cluster expression across species, with the piRNA in the lower part and the RNA expression per kb +1 in the upper part. Clusters are coloured by timing of expression annotated for the overlapping Yu2021 piRNA clusters. (C) RNA versus piRNA log2FC in distinct contrasts of differential expression analyses, showing that most significant changes occur only at piRNA level, while few occur at RNA or both piRNA and RNA levels. (D) Proportion of pachytene Yu2021 piRNA clusters bound by A-MYB (left) or TCFL5 (right) at their promoter (TSS +/-500bp), stratified by conservation level (there were no clusters overlapping Yu2021 and conserved in only one species, 6 conserved in two species and 57 conserved in all three) based on the overlap to predicted piRNA clusters expressed in *Mus musculus.* Fisher’s exact test (conserved in two versus three species): p = 1 for both factors.

**Supplementary figure 5.**
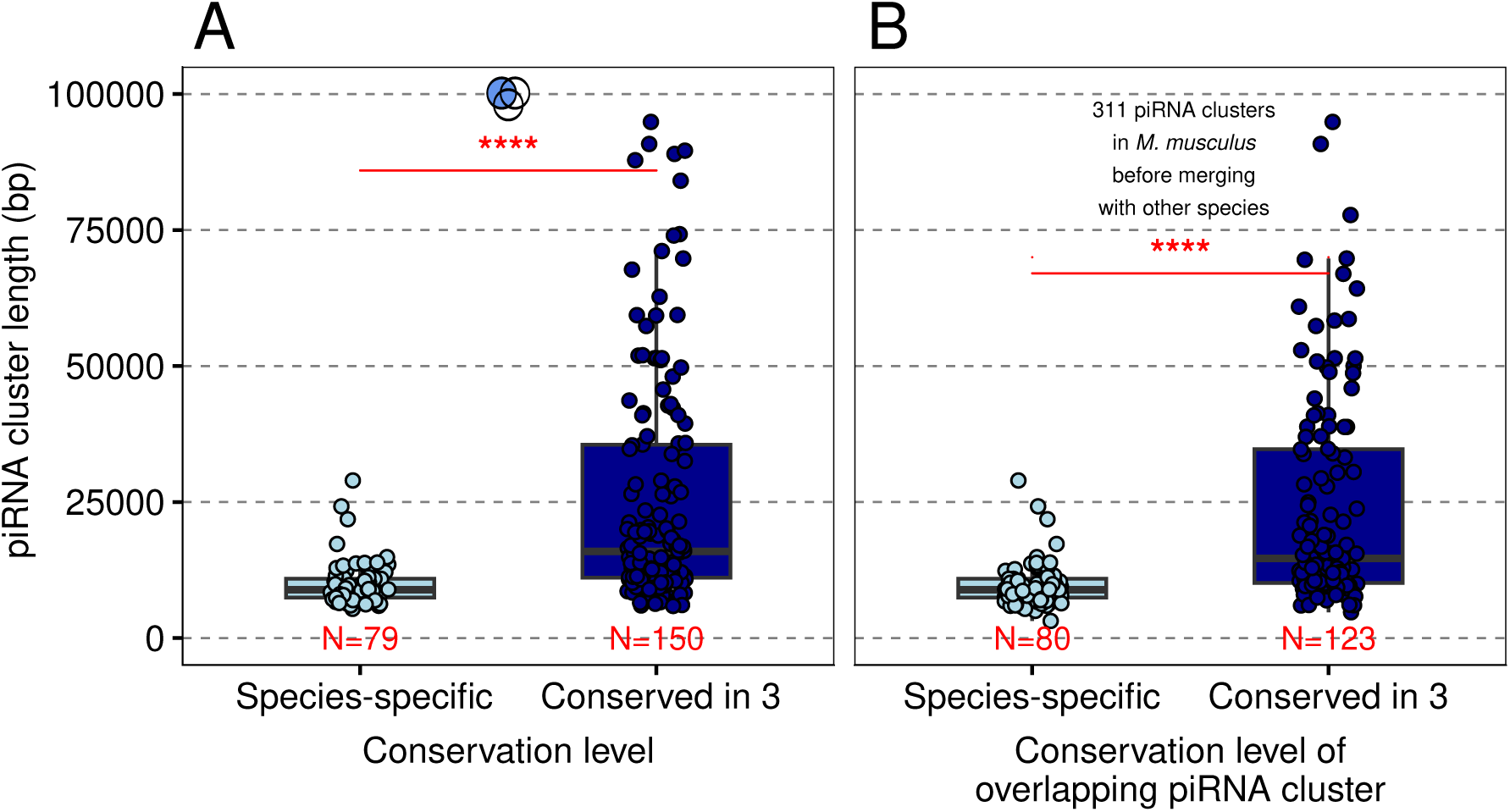
Length of predicted piRNA clusters increases with conservation. (A) Length of final species-specific (N=79) and conserved (N=150) piRNA clusters predicted in *Mus musculus*. (B) Length of the 311 piRNA clusters predicted in *Mus musculus* before merging them with the other species and overlapping them to species-specific (N=80) or conserved (N=123) piRNA clusters from the final set. Significance values are from Wilcoxon rank sum (non-paired) test. Significance codes: *: p<0.05, **: p<0.01, ***: p<0.001, ****: p<0.0001.

**Supplementary figure 6.**
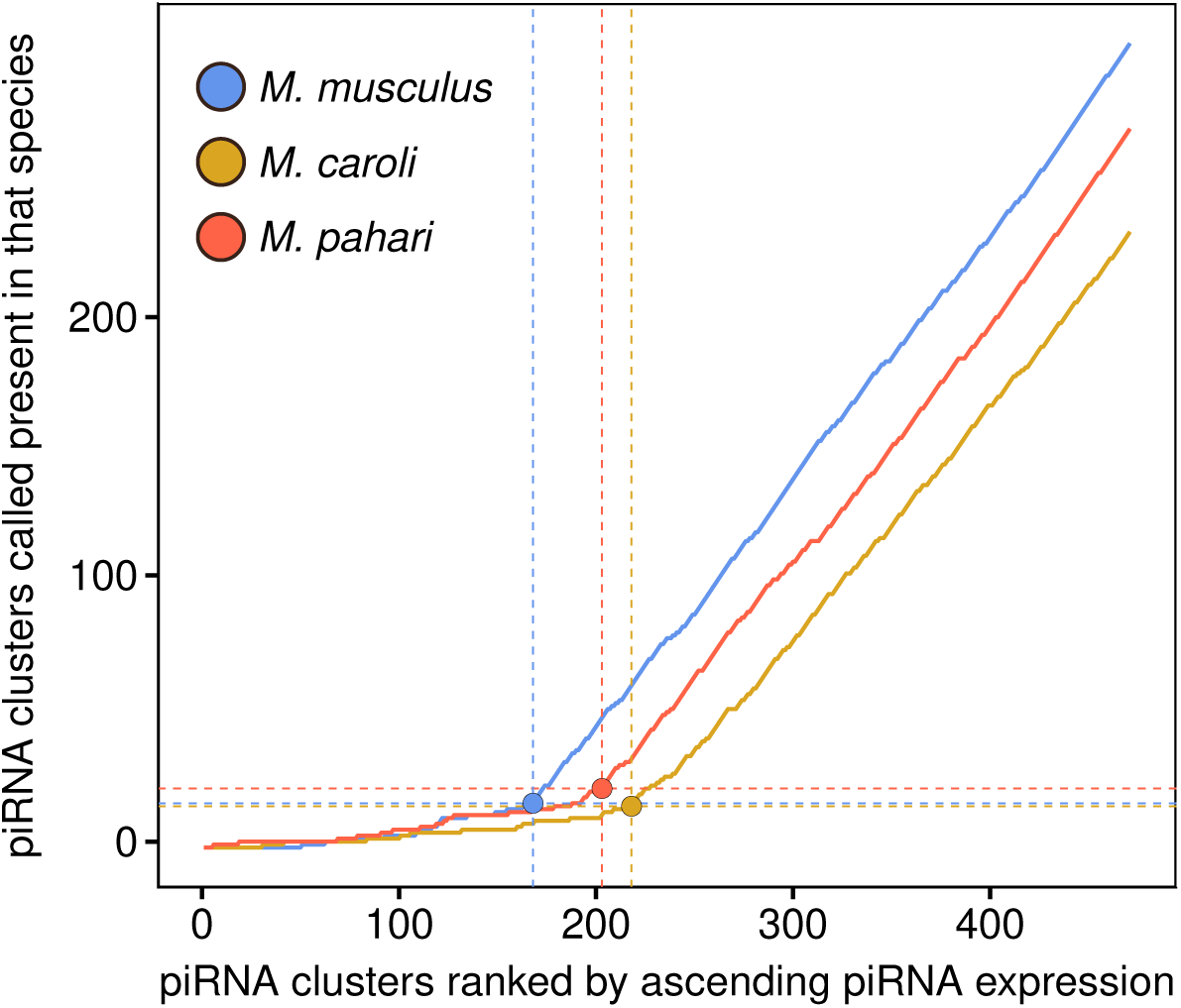
Determination of per-species expression thresholds for defining piRNA-producing loci. Loci ranked by ascending piRNA expression in each species (x axis) against the cumulative number of loci called as piRNA clusters by proTRAC in that species (y axis). Filled circles and dashed lines mark the inflexion point used as the per-species expression threshold to define piRNA-producing loci (see Methods).

**Supplementary figure 7.**
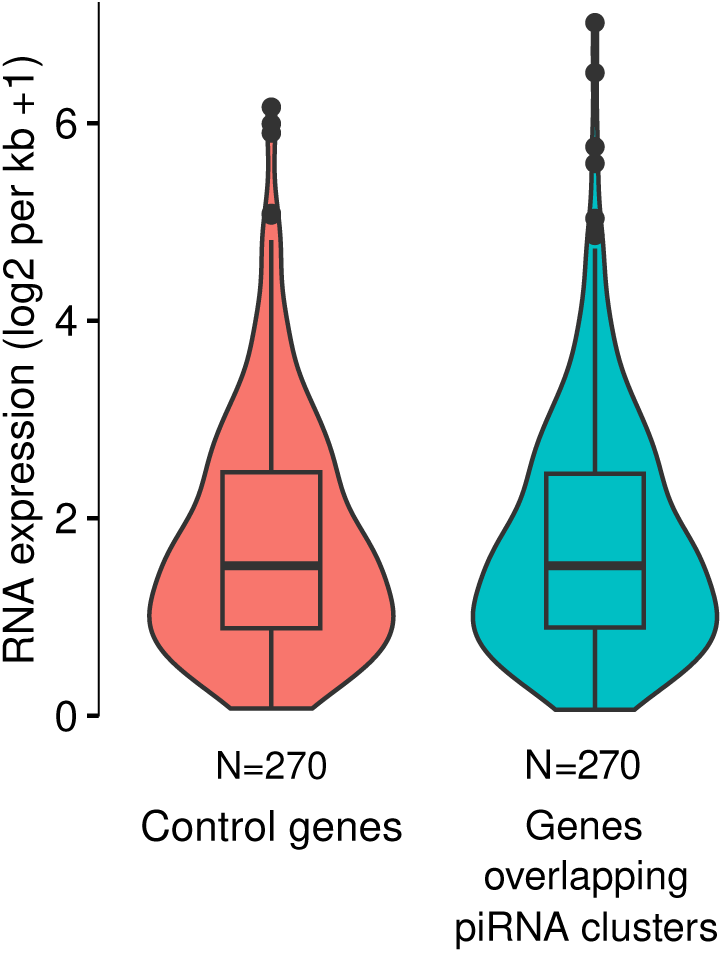
Distribution of transcription level of genes overlapping piRNA clusters and the piRNA cluster expression-matched control genes. Violin plots of RNA expression (log2 per kb +1) in genes overlapping piRNA clusters and in the expression-matched control genes selected as described in Methods.

## SUPPLEMENTARY TABLES

**Supplementary table 1. Sample information and sequencing statistics.**

**Supplementary table 2. GRCm39/mm39 coordinates of predicted piRNA clusters in each species before merging them.**

-A. piRNA clusters predicted in M. musculus.

-B. piRNA clusters predicted in M. caroli.

-C. piRNA clusters predicted in M. pahari.

**Supplementary table 3. Coordinates of final predicted piRNA clusters in each species assembly.**

-A. Coordinates of the 215 Yu2021 piRNA clusters. Note that piRNA clusters with NA values are those not present in all species after conversion with ENSEMBL Compara.

-B. Coordinates of the 437 predicted piRNA clusters.

**Supplementary table 4. Results of the piRNA cluster – TE overlap permutation test.**

**Supplementary table 5. TE coverage in piRNA clusters and control genes in M. musculus.**

-A. TE coverage in piRNA cluster expression-matched control genes.

-B. TE coverage in Yu2021 piRNA clusters.

-C. TE coverage in predicted piRNA clusters.

**Supplementary table 6. UTR length of control genes and genes overlapping piRNA clusters.**

-A. UTR length in piRNA cluster expression-matched protein-coding control genes.

-B. UTR length in protein-coding genes overlapping Yu2021 piRNA clusters.

-C. UTR length in protein-coding genes overlapping predicted piRNA clusters.

-D. UTR length of protein-coding genes overlapping species-specific piRNA clusters and their non piRNA-producing orthologous genes.

**Supplementary table 7. TE coverage in 3’UTRs of control genes and genes overlapping piRNA clusters.**

-A. TE coverage in 3’UTRs of piRNA cluster expression-matched protein-coding control genes.

-B. TE coverage in 3’ UTRs of protein-coding genes overlapping Yu2021 piRNA clusters.

-C. TE coverage in 3’UTRs of protein-coding genes overlapping predicted piRNA clusters.

**Supplementary table 8. AREscore and length-normalized AREscore in UTRs of control genes and genes overlapping piRNA clusters.**

-A. AREscore in UTRs of piRNA cluster expression-matched protein-coding control genes.

-B. AREscore in UTRs of protein-coding genes overlapping Yu2021 piRNA clusters.

-C. AREscore in UTRs of protein-coding genes overlapping predicted piRNA clusters.

**Supplementary table 9. Transcriptional readthrough in control genes and genes overlapping piRNA clusters.**

-A. Transcriptional readthrough in piRNA cluster expression-matched control genes.

-B. Transcriptional readthrough in genes overlapping Yu2021 piRNA clusters.

-C. Transcriptional readthrough in genes overlapping predicted piRNA clusters.

**Supplementary table 10. TE coverage in downstream-of-gene (DoG) regions of control genes and genes overlapping piRNA clusters.**

-A. TE coverage in DoG regions of piRNA cluster expression-matched control genes.

-B. TE coverage in DoG regions of genes overlapping Yu2021 piRNA clusters.

-C. TE coverage in DoG regions of genes overlapping predicted piRNA clusters.

-D. TE coverage in DoG regions in all genes with transcriptional readthrough.

**Supplementary table 11.**

**Supplementary table 12. Small RNA-seq expression in 437 predicted piRNA clusters.**

-A. Raw counts

-B. DEseq2-normalized counts

-C. Differentially expressed piRNA clusters

**Supplementary table 13. Total RNA-seq expression in 437 predicted piRNA clusters.**

-A. Raw counts

-B. DEseq2-normalized counts

-C. Differentially expressed piRNA clusters

**Supplementary table 14. Small RNA-seq expression in 184 Yu2021 piRNA clusters.**

-A. Raw counts

-B. DEseq2-normalized counts

-C. Differentially expressed piRNA clusters

**Supplementary table 15. Total RNA-seq expression in 184 Yu2021 piRNA clusters**

-A. Raw counts

-B. DEseq2-normalized counts

-C. Differentially expressed piRNA clusters

**Supplementary table 16. Relationship between piRNA expression in a focal species and piRNA production in a target species. piRNA expression in one species predicts piRNA production at the orthologous locus in another species.**

Odds ratio (95% confidence interval) that the locus is a piRNA-producing locus in the target species, for each two-fold increase in piRNA expression in the focal species (binomial logistic regression).

**Supplementary table 17. Ordinal logistic regression of piRNA cluster conservation on piRNA expression and other features in mouse.**

Ordinal logistic regressions modelling piRNA cluster conservation levels against piRNA expression and other piRNA cluster features in the three mouse species. Model A uses piRNA expression as predictor of conservation levels; model B includes piRNA expression and precursor RNA expression; model C uses piRNA cluster width and piRNA expression not normalized by width; model D includes piRNA expression and the presence of transcriptional readthrough; and model E uses all the previous features to predict conservation levels. Focal species is included as a predictor to account for piRNA clusters appearing multiple times. Conservation levels are species-specific (expressed only in one species), close (expressed in M. musculus and M. caroli), ambiguous (expressed in M. pahari and one of the other species) and conserved piRNA clusters (expressed in all three species). The ANOVA tests compare each model with the model A. Expression is normalised by cluster width (expression per kb) except for when width is included (abundance). RNA, piRNA expression and piRNA cluster width are log2-transformed.

**Supplementary table 18. Coordinates and information of 182 human Ozata2020 piRNA clusters.**

**Supplementary table 19. Small RNA-seq expression in 182 human Ozata2020 piRNA clusters.**

-A. Raw counts

-B. DEseq2-normalized counts

**Supplementary table 20. Total RNA-seq expression in 182 human Ozata2020 piRNA clusters**.

-A. Raw counts

-B. DEseq2-normalized counts

**Supplementary table 21. UTR length of protein-coding genes overlapping 182 human Ozata2020 piRNA clusters.**

**Supplementary table 22. Transcriptional readthrough in genes overlapping 182 human Ozata2020 piRNA clusters.**

**Supplementary table 23. Ordinal logistic regression of piRNA cluster conservation on piRNA expression and other cluster features in human.**

Ordinal logistic regressions modelling piRNA cluster conservation levels against piRNA expression and other piRNA cluster features in the human piRNA clusters. Model A uses piRNA expression as predictor of conservation levels; model B includes piRNA expression and transcriptional level; model C includes piRNA expression and the presence of transcriptional readthrough; and model D uses all the previous features to predict conservation levels. Conservation levels are species-specific (expressed only in human), close (expressed in human and macaque), ambiguous (expressed in human and marmoset) and primate-conserved piRNA clusters (expressed in all three species). The ANOVA tests compare each model with the model A. Expression is normalized by piRNA cluster width (per kb) and log2- transformed.

**Supplementary table 24. piRNA cluster expression-matched control genes.**

